# Short-term Pre-operative Methionine Restriction Induces Browning of Perivascular Adipose Tissue and Improves Vein Graft Remodeling in Mice

**DOI:** 10.1101/2023.11.02.565269

**Authors:** Peter Kip, Thijs J. Sluiter, Michael R. MacArthur, Ming Tao, Jonathan Jung, Sarah J. Mitchell, Sander Kooijman, Nicky Kruit, Josh Gorham, Jonathan G. Seidman, Paul H.A. Quax, Masanori Aikawa, C. Keith Ozaki, James R Mitchell, Margreet R. de Vries

## Abstract

Short-term preoperative methionine restriction (MetR) shows promise as a translatable strategy to modulate the body’s response to surgical injury. Its application, however, to improve post-interventional vascular remodeling remains underexplored. Here, we find that MetR protects from arterial intimal hyperplasia in a focal stenosis model and adverse vascular remodeling after vein graft surgery. RNA sequencing reveals that MetR enhances the brown adipose tissue phenotype in arterial perivascular adipose tissue (PVAT) and induces it in venous PVAT. Specifically, PPAR-α was highly upregulated in PVAT-adipocytes. Furthermore, MetR dampens the post-operative pro-inflammatory response to surgery in PVAT-macrophages *in vivo* and *in vitro*. This study shows for the first time that the detrimental effects of dysfunctional PVAT on vascular remodeling can be reversed by MetR, and identifies pathways involved in browning of PVAT. Furthermore, we demonstrate the potential of short-term pre-operative MetR as a simple intervention to ameliorate vascular remodeling after vascular surgery.

**Graphical Abstract:** 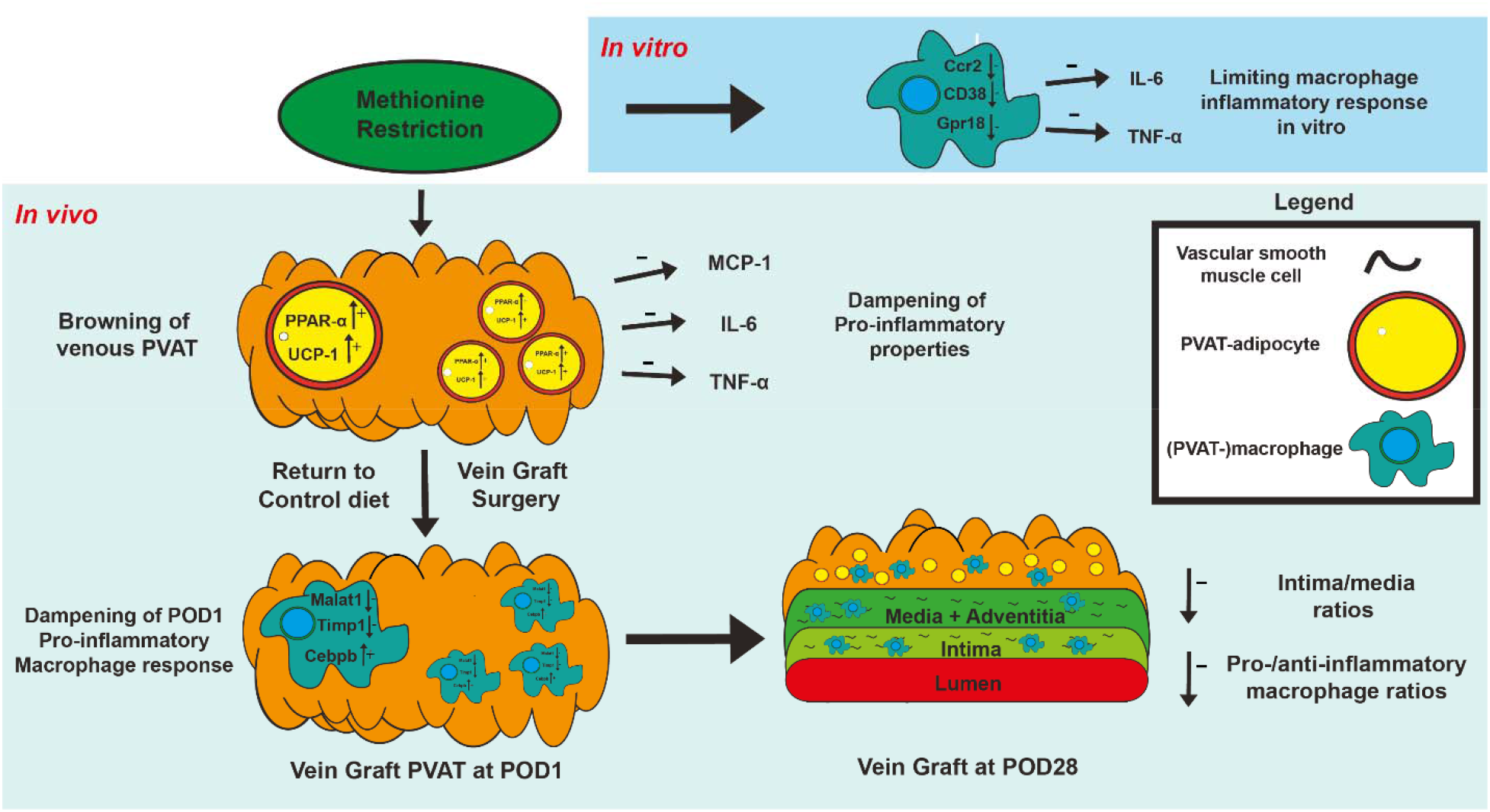

## Introduction

Revascularization surgery remains a mainstay in the treatment of arterial occlusive disease^1^, however these interventions are hampered by high failure rates which frequently originate from intimal hyperplasia (IH) and adverse vascular remodeling.^2^ This pathophysiological response to surgical injury is defined by initial endothelial dysfunction and leukocyte transmigration^2, 3^ followed by vascular smooth muscle cell (VSMC) migration and proliferation, ultimately occluding the artery or vein graft.^2, 4^ Despite decades of research, therapies to abrogate adverse vascular remodeling responses following an intervention remain limited.

Perivascular adipose tissue (PVAT) surrounding blood vessels functions as a paracrine organ with the potential for impacting post-interventional vascular remodeling.^5, 6^ Interestingly, preclinical work suggests inflamed PVAT driven by obesity can accelerate IH.^6^ For example, high-fat diet (HFD) in mice increases the secretion of interleukin (IL)-6, IL-8 and C-C motif ligand 2 (CCL2) from perivascular adipocytes.^7^ After carotid wire injury, cohorts in which inflamed adipose tissue was induced by HFD had exacerbated IH.^8, 9^

Surgical trauma to the fat itself also alters its local phenotype and can even yield a systemic response in distal adipose depots.^10^ In mice, unilateral surgical trauma to inguinal adipose tissue induces local and distal browning^11^, but whether this could be beneficial in post-operative recovery is unknown. In this same model, lowering of the high-fat content of the diet for the 3 weeks leading up to the surgical injury not only reduced baseline levels of CCL2 and tissue necrosis factor-α (TNF-α), but also dampened the inflammatory response of the fat to surgical trauma. At post-op day (POD)-1, local levels of both IL-1β and TNF-α were reduced compared to mice who received a preoperative HFD.^10^

This concept of utilizing short-term dietary interventions before surgery to precondition the body stands as an emerging approach to enhance surgical outcomes.^12, 13^ Dietary restriction (DR), defined as restriction of either total calories, macronutrients, or specific essential amino acids, represents one such dietary preconditioning approach.^13, 14^ Efficacy of these short-term diets has been shown in a wide range of preclinical surgical models, including renal^12, 15, 16^, hepatic^12, 17, 18^ and vascular injury models.^19^ Recently, we reported^20^ that short-term dietary protein restriction limits adverse vascular remodeling in a murine venous bypass graft model via increased expression of cystathionine y-lyase. This enzyme is part of the transsulfuration pathway, and activation results in increased endogenous production of the gaseous vasodilatory and anti-inflammatory molecule hydrogen sulfide.^18^ Hydrogen sulfide in turn was able to function as a DR-mimetic when local levels were exogenously increased during vein graft surgery via a locally applied gel, thereby limiting adverse vein graft remodeling.^21^

Methionine restriction (MetR) is a DR regimen in which dietary sulfur amino acid (methionine and cysteine) content, but not overall calorie intake, is reduced. In rodents, MetR has pleiotropic beneficial effects on markers of cardiometabolic health^22^ and lifespan^23^ likely via effects on adipose tissue^24^ and energy metabolism^25^, while specifically in surgical models MetR improves outcome of femoral ligation by increasing angiogenic potential^26^ and preserves wound healing.^27^ In humans, MetR delivered for up to 16 weeks as a semi-synthetic diet is feasible and increases fat oxidation and reduces intrahepatic lipid content.^28^

Here we tested the hypothesis that short-term pre-operative MetR can attenuate adverse vascular remodeling in models of arterial injury and vein graft surgery specifically via interplay with PVAT, with implications for dietary preconditioning in human vascular injury. Mechanistically, we evaluated changes in PVAT gene expression prior to and after surgery and their modulation by MetR by bulk and nuclear RNA sequencing.

## Methods

### Experimental animals

All animal experiments were approved by the appropriate Brigham and Women’s Hospital Institutional Animal Care and Use Committee and in accordance with the NIH recommendations for care. All surgical experiments were performed on C57BL/6 mice (male, 14-16 weeks old, Stock No: 380050, Jackson Laboratory). Male mice were used in this study due to our earlier findings that estrogen inhibits neointima formation,^29^ and previous work showing that MetR responses are highly sexually dimorphic, with females showing resistance to MetR metabolic benefits.^30, 31^ We recognize this as a limitation and future research should optimize a female-specific MetR paradigm to allow replication of these findings in female mice. Mice were housed 4-5 per cage and maintained on a 12-hour light-dark cycle at 22°C with 30-50% humidity. The animal facility was specific pathogen-free (SPF) as confirmed by quarterly sentinel animal testing.

### Dietary Intervention

All mice (aged 10-12 weeks old) were started on a 3-week 60% fat (by calories), 0% cysteine diet (Research Diets, A18013001) [Control] which contained standard levels of methionine (0.6%, 2.6% of total protein). After 3 weeks of Control diet, 1 cohort was switched to a methionine restriction (MetR) diet containing 60% fat, 0% cysteine and 0.07% methionine, 0.3% of total protein) [Research Diets, A18022602]. After 1 week of MetR or continued Control diet, mice were either harvested for caval vein/baseline studies or underwent a surgical intervention. Immediately post-operatively, all mice were switched back to the Control diet and tissue was harvested at either POD1 or POD28. (**Fig. 1A**)

**Figure 1.**
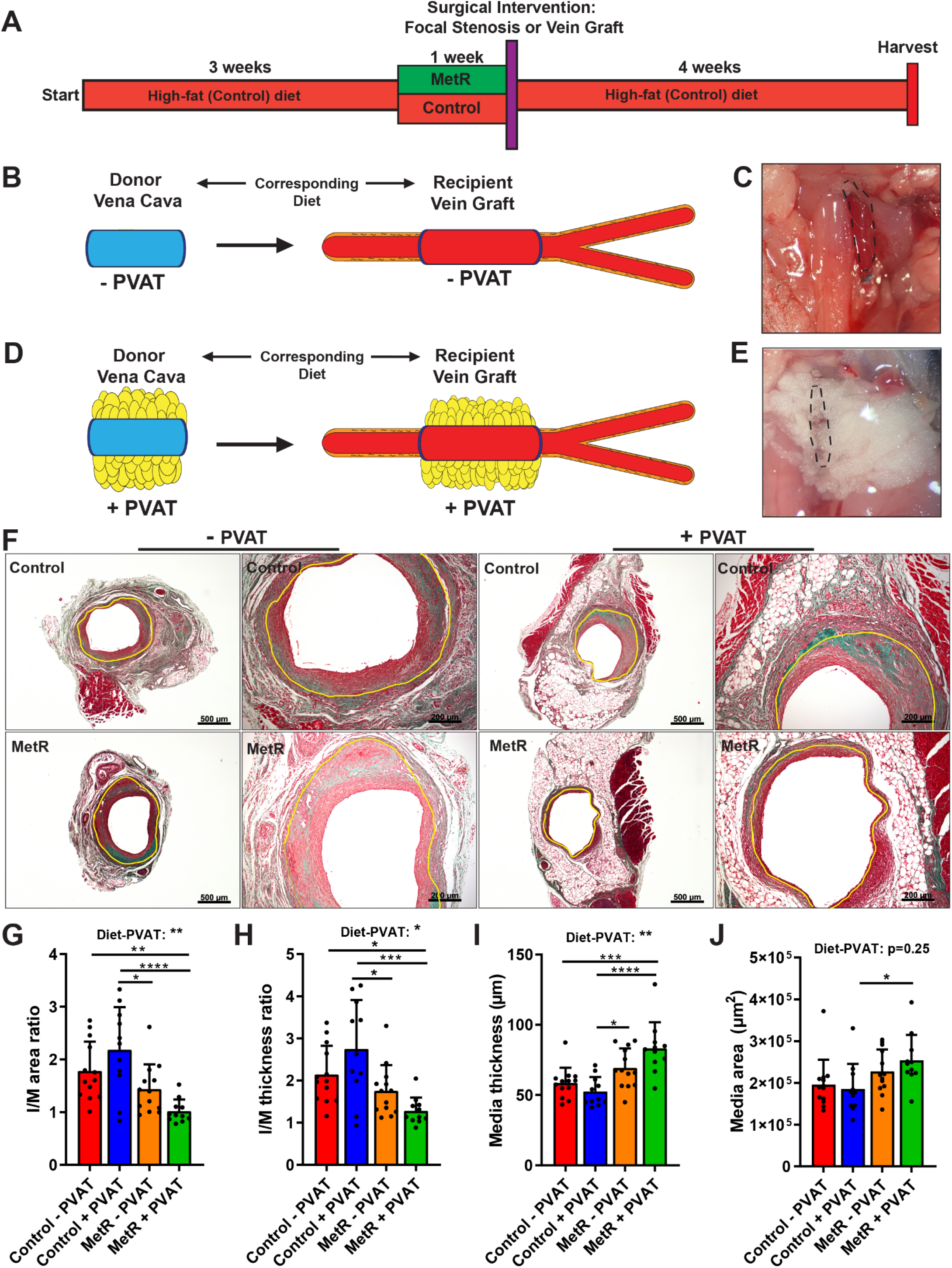
Protection from Adverse Vein Graft Remodeling via Short-term Methionine Restriction is Perivascular Adipose Tissue Dependent. **A:** Schematic overview of dietary intervention. **B-E:** schematic and in-situ images of vena cava/vein graft +-PVAT. **B,C**: stripping of vena cava PVAT results in vein graft lacking PVAT. **D,E**: vena cava with PVAT and consecutive vein graft with PVAT intact. **F**: Images of vein grafts at POD28 after Masson-trichrome staining. Control-fed and MetR, no PVAT or PVAT. Scale bars = 200µm or 500µm as indicated. **G-J**: histomorphometric analysis of POD28 vein grafts. **G**: I/M area ratio. **H**: I/M thickness ratio. **I:** Media thickness. **J**: Media area. All statistical testing was done via two-way ANOVA with Tukey’s multiple comparisons test unless otherwise indicated, n=11-13/group. * P<0.05, ** P<0.01, *** P<0.001, **** P<0.0001

### Vein Graft Surgery

Vein graft surgery was performed as described previously.^32^ In brief, mice were anesthetized with isoflurane for the duration of the procedure. Shortly before the start of the recipient surgical procedure, the thoracic caval vein of a donor mouse was harvested and placed in sterile 0.9% NaCl supplemented with heparin (100UI/mL). In the recipient, neck region fur was removed, and a neckline incision was performed. The right common carotid artery (RCCA) was dissected from its surrounding soft tissues and 2 8-0 nylon sutures were tied in the middle, approximately 1mm apart. RCCA was then cut between the two sutures to facilitate an end-to-end anastomosis. The proximal and distal RCCA was then everted over an autoclavable nylon cuff (Portex) of approximately 2mm while clamped with vascular clamps. The everted carotid walls were secured with an 8-0 nylon suture. Next the donor caval vein was sleeved between both RCCA ends, and an end-to-end anastomosis was created with 8-0 nylon sutures. The distal followed by the proximal vascular clamp was released to restore blood flow. The incision was closed with 6-0 Vicryl sutures. Post-operatively animals received warm Ringer’s lactate solution (0.5mL, subcutaneous) and buprenorphine (0.1mg/kg, subcutaneous).

### Focal Stenosis Creation

A focal stenosis was created as described previously^33^ to generate an arterial intimal hyperplastic response. Anesthesia was induced via 4% isoflurane and maintained via 2-3% isoflurane in a nose cone for the duration of the procedure. The RCCA was dissected from its surrounding tissue. A 35-gauge blunt needle mandrel was then placed longitudinally along the RCCA and tied with a 9-0 nylon suture approximately 2-2.5mm proximal to the bifurcation. After removal of the needle mandrel, the skin was closed with a 6-0 Vicryl suture. Post-operatively mice received warm Ringer’s lactate solution (0.5mL, subcutaneous) and buprenorphine (0.1mg/kg, subcutaneous).

### Vein Graft PVAT Manipulation

In routine rodent vein graft surgery as well as in our first cohort of vein graft dietary intervention experiments, the donor caval vein was partially trimmed of its surrounding PVAT to facilitate end-to-end anastomosis creation in the recipient. In a follow-up cohort of C57BL/6, we either completely stripped the donor caval vein of its surrounding PVAT or left all PVAT intact. The donor caval vein (with/without PVAT) was then transplanted into a recipient RCCA on a corresponding diet (Control or MetR), to create a vein graft with PVAT intact, or a vein graft lacking PVAT. This resulted in 4 separate groups evaluating the interplay of diet and the presence of PVAT around the vein graft: Control – PVAT, Control + PVAT, MetR – PVAT, MetR + PVAT. Stripped donor caval vein PVAT was snap frozen in liquid nitrogen and then stored at -80°C for subsequent analyses.

### Vein graft/RCCA POD28 Harvest

Under isoflurane anaesthesia, mice were euthanized via exsanguination, followed by insertion of a 21G needle in the left ventricle. Whole-body perfusion was performed with Ringer’s lactate solution for 3 minutes, and followed by 3 minutes of perfusion-fixation with 10% formalin. The graft/RCCA was excised en-bloc via a midline neck incision and transferred to a 10% formalin (in PBS) solution for 24 hours. After 24 hours the tissue was transferred to a 70% ethanol solution for further processing as described below.

### Baseline Studies

To study effects of diet without surgery, baseline data was obtained from mice fed an identical dietary intervention (3-week control diet followed by 1-week MetR or Control) and harvested after 1 week of MetR/Control. All tissue was collected on dry ice and snap frozen in liquid nitrogen, before storage at -80°C. For baseline harvest, mice were first anesthetized, and a cardiac puncture was performed to collect 1mL of whole blood in a 1.5mL ethylenediaminetetraacetic acid (EDTA)-coated Eppendorf for downstream blood analysis. A thoracotomy was performed and caval vein PVAT was carefully dissected from the vessel wall with forceps. The caval vein was then harvested and caval vein wall and PVAT were stored separately. Lungs and heart were removed and thoracic aorta PVAT was carefully separated from the vessel wall, followed by harvest of the aortic wall. Both aortic and caval PVAT were stored separately.

### Vein Graft POD1 Harvest

For vein graft POD1 harvest, after mice were anesthetized, the neck suture was removed and the surgical field from the previous day was opened. Vein graft patency was first ensured visually. In one cohort of mice, up to 1mL whole blood was collected via cardiac puncture in an EDTA-coated Eppendorf for downstream analysis, and then the vein graft was removed for histology as follows. A thoracotomy was performed, followed by whole-body perfusion via the left ventricle with ringer’s lactate solution. A 21G syringe containing O.C.T. (Tissue-Tek, # 25608-930) was carefully inserted in the brachiocephalic trunk oriented towards the RCCA. Next, O.C.T. was slowly released into the brachiocephalic trunk until the vein graft started to dilate. After placing an 8/0 suture around the distal cuff, it was tightened followed by a second suture around the proximal cuff. The now dilated vein graft was removed en-bloc and placed in a mold filled with O.C.T., then placed on dry ice and stored in -80°C. In a second cohort of mice, the PVAT surrounding the vein graft was carefully removed using forceps and collected in a 1.5mL Eppendorf on dry ice. Next, the vein graft wall itself was taken out and collected in a separate Eppendorf on dry ice. Both PVAT and vein graft were then snap frozen in liquid nitrogen and stored at -80°C.

### Vein Graft/RCCA Histology

POD28 Vein grafts and RCCA (at POD28 after focal stenosis surgery) were harvested, embedded in paraffin, and cut in 5µm sections by microtome, then mounted on slides. Sections were collected at regular intervals of 200µm, starting from the proximal cuff until 1000µm post proximal cuff. Focal stenosis arteries were cut at regular intervals of 400µm, starting at 400µm proximal from the focal stenosis, until 2800µm proximal from the stenosis. For histomorphometric analysis, a Masson-Trichrome staining was performed: after deparaffinization to 95% ethanol, slides were immersed in 5% picric acid (in 95% ethanol) for 3 minutes, followed by a 3-minute stain in working Harris Hematoxylin Solution (Fisher 213 Scientific, cat# 245-678). After a brief tap water wash, slides were stained with 1% Biebrich Scarlet in 1% acetic acid (Fisher Scientific, cat# A38S-500) for 3 minutes, followed by a quick rinse in distilled water. Slides were then stained for 1 minute in 5% Phosphomolybdic/Phosphotungstic acid solution and immediately transferred to 2.5% light green SF yellowish in 2.5% acetic acid (Fisher Scientific, cat# A38S-500) for 4 minutes. Followed by a quick rinse in distilled water and a 2-minute rinse in 1% acetic acid solution (Fisher Scientific, cat# A38S-500). After dehydration with Xylene, slides were covered with a cover glass employing Permount (Electron Microscopy Science, cat# 17986-05). Brightfield images of vein graft and carotid artery cross-sections were taken with a Zeiss Axio A1 microscope (Carl Zeiss). Histomorphometric analysis was performed using Image J 1.51p (Java 1.8.0_66) (see below).

### Histomorphometric Analysis

For vein graft histomorphometric analysis, images of cross sections taken at 200µm, 400µm, 600µm, 800µm, 1000µm and 1200µm post-cuff were uploaded in ImageJ. Per distance, 1 cross section was analyzed. Area and perimeter of lumen, internal elastic lamina and adventitial border were measured in µm^2^/µm. Next, lumen area, intimal area, intimal thickness, media+adventitia (M) area, M thickness, intimal/media+adventitia (I/M) area ratio and I/M thickness ratio were calculated as described previously.^34^ 200µm-1000µm cross sections were next averaged into a per-vein graft histomorphometric endpoint. For focal stenosis histomorphometric analysis, RCCA cross section images taken at 400µm, 800µm, 1200µm, 1600µm and 2000µm proximal from the focal stenosis were uploaded in ImageJ. Lumen area and perimeter, internal elastic lamina area and perimeter, external elastic lamina area and perimeter were measured. Next, lumen area, intimal area and thickness, medial area and thickness and I/M area and thickness ratios were calculated. Collagen measurements were performed via the color deconvolution function in ImageJ, the resulting split green area was measured via pixel-threshold and normalized to total intimal/M and total vein graft area in %.

### Immunohistochemistry

First, vein graft slides were incubated for 30 minutes at 60°C in a vacuum oven, followed by immediate deparaffinization. Next, antigen retrieval was performed for 30 minutes at 97°C in Citrate buffer (pH 6.0) [Abcam, ab93678]. Then, slides were pre-incubated with 10% goat serum (Life Technologies, 50062Z) in PBS with 0.3M Glycine (Aijomoto, R015N0080039) for 1hr at room temperature (RT). Slides were then incubated with primary antibodies. For VSMC+KI-67 double-staining: ACTA2 (mouse anti-mouse, Abcam, ab7817, 1:800) and Ki-67 (rabbit anti-mouse, Abcam, ab16667, 1:100). For M1/M2 macrophage staining, 1 vein graft slide containing 2-4 cross sections was double stained with the general macrophage marker Mac-3 (rat anti-mouse, Fisher Scientific, B550292, 1:600) and either iNOS (rabbit anti-mouse, abcam, ab3523, 1:100) for M1, or CD206 (rabbit anti-mouse, abcam, ab64693, 1:800) for M2. Slides with primary antibodies were incubated overnight at 4°C. Slides were then washed in PBS + tween (PBST) and incubated in secondary antibody for 2hrs at RT. For ACTA2 + Ki-67 double staining, slides were incubated with Alexa Fluor 647 (goat anti-mouse, A-32728) and Alexa Flour 568 (goat anti-rabbit, A-11011) at 1:600. For M1/M2 staining, slides were incubated with Alexa Fluor 568 (goat anti-rabbit, A-11011) and Alexa Fluor 647 (goat anti-rat, A-21247) at 1:600. After secondary antibody incubation, slides were washed in PBST and mounted with DAPI (Vector, CB-1000) or stained with Hoechst staining solution and mounted with anti-fade as indicated. For immunohistochemistry (UCP1 / PPAR-α staining) slides were deparaffinized, endogenous peroxidase was blocked with 0.3% H_2_O_2_ in PBS for 15 minutes (only for UCP1), followed by antigen retrieval in Tris EDTA buffer (DAKO Blue) for 10 minutes at 95°C (UCP1) or 98°C (PPAR-α). Then, slides were pre-incubated with blocking solution (PBS + 5% albumin (PBSA) for UCP1, or 2.5% PBSA + 1% goat serum for PPAR-α), after which slides were incubated overnight in a humidity chamber with primary antibodies: UCP1 (rabbit anti-mouse, Abcam, ab155117, 1:400), PPAR-α (rabbit anti-mouse, ThermoFisher, BS-3614R, 1:200). Slides were then washed in PBST, 25 minutes incubated in 0.3% H_2_O_2_ in PBS and again washed in PBST (only for PPAR-α). Secondary antibody (Envision anti-rabbit) was applied and slides were incubated for 45 minutes at RT in the humidity chamber. After incubation, slides were washed with PBST, DAB solution (1mL of substrate + 1 drop of chromogen) added for 7 minutes, washed with diH_2_O, immersed in Hematoxylin staining during 10 seconds and rinsed with running tap water. Slides were then dehydrated and mounted with Pertex and coverslips.

### Immunohistochemical analysis

For fluorescent IHC, 20x images were taken with a Laser Scanning Confocal Microscope (Zeiss LSM800) and automatically stitched. For non-fluorescent IHC, 10x brightfield images were taken with a Zeiss Axio A1 microscope (Carl Zeiss) and stitched with Adobe Photoshop. Lumen area, intimal area, media area and PVAT area were defined and measured in ImageJ, based on the corresponding Masson-trichrome histology picture. For VSMC + Ki-67 double staining analysis, 1 cross section per vein graft was analyzed (600 or 800µm). Based on Hoechst/DAPI staining, the lumen, intima and media area was measured in mm^2^. The ACTA2 positive fluorescent (647nm) channel was analyzed for total ACTA2 positive pixels via color-threshold and then normalized to its respective vein graft layer (intima, media or total vein graft) in %. A cell positive for both ACTA2 and Ki-67 was regarded as a “proliferating VSMC”. Total proliferating VSMCs per vein graft layer were then counted and normalized to cells/mm^2^. Total ACTA2 positive cells were counted per vein graft layer and normalized to VSMC/mm^2^ per vein graft layer. For analysis of macrophage polarization, one slide per vein graft (at 800µm) containing 2-4 cross sections was stained with Mac-3+iNOS or Mac-3+CD206. Lumen area, intimal area, media area and PVAT area were measured in mm^2^.

All cells positive for Mac-3 were determined macrophages (M_□_). Pro-inflammatory macrophage were cells positive for Mac-3 and iNOS. Anti-inflammatory macrophage were cells positive for Mac-3 and CD206. In each cross section the total number of M_□_ and Pro/-anti-inflammatory - macrophages per vein graft layer was counted and normalized to mm^2^. Per vein graft layer, Pro- and anti-inflammatory macrophages (in count per mm^2^) were normalized as a percentage of the total M -macrophages present in that layer (also in count/mm^2^). The resulting percentages were then used to calculate a Pro-/anti-inflammatory macrophage ratio.

For immunohistochemical analysis (UCP1 / PPAR-α staining / adipocyte size) slides were automatically scanned used a 3D histec slide scanner and images were uploaded in ImageJ. For the UCP1 and PPAR-α staining, the positive PVAT-area was divided by the total PVAT area, which was manually selected. The positive area was quantified via intensity-based processing: the image was changed to HSB-brightness stack, in order to highlight the UCP1 positive area. Via thresholding, these areas were automatically selected, calculated and divided over the total PVAT area. To calculate the size of the adipocytes, Masson-Trichrome stained slides were used. First, the PVAT-area was manually selected, after which customized scripts were used to automatically determine the adipocyte size. In brief, images were converted to grayscale and a threshold was set to create a black and white image. The image was then segmented using the watershed option in ImageJ to identify distinct adipocytes. The size was measured for all segmented adipocytes and plotted in a size-distribution graph.

### Cell culture under MetR conditions

MetR medium was prepared by titrating 0% MetR medium (complete DMEM, ThermoFisher, 21013-024, 10% FCS) with 100% MetR medium (complete DMEM without L-methionine and L-cysteine, ThermoFisher, 11960-044, 10% FCS). For example, 1 part 0% MetR medium was mixed with 3 parts 100% MetR medium to create 75% MetR medium.

### Bone marrow-derived macrophages

Bone marrow-derived macrophages (BMDM) were harvested by removing the proximal and distal end of both femur and tibia of multiple mice. Bones were flushed with cold PBS, collected through a cell strainer (70µm) and spun down at 1000 rpm for 10 minutes. The pellet was washed with PBS and resuspended in 3 mL of ACK lysis buffer followed by 3 minutes of incubation on ice. Lysis was stopped by adding 7 mL of RPMI/25% FCS, after which the suspension was washed twice with PBS. Cells were counted using a Hemacytometer and seeded at 8×10^6^ cells / dish (Falcon 100×15mm) in 10 mL of medium (RPMI/25% FCS) with 20µg/ml M-CSF. Medium was changed 7 days after seeding and after an additional 3 days cells were seeded at 300.000 cells / well in a 24-well plates and left to adhere overnight. Cells were washed and control or MetR medium supplemented with 10 ng/mL LPS (Sigma, K235) was added. After 24 hours, medium was transferred to an Eppendorf and stored at -80°C. Trizol was added to the cells and the plate was stored at -20°C for RNA extraction.

### 3T3-L1 adipocytes

Murine 3T3-L1 preadipocytes differentiated into mature adipocytes by first incubating confluent cells in induction-medium (DMEM GlutaMAX™, Gibco®, 10% FCS) containing 1.6μM insulin, 0.5mM 1-methyl-3-isobutylxanthine (IBMX), 0.25μM dexamethasone. All medium contained 100U/mL penicillin/streptomycin. After 2 days, cells were washed and differentiation-medium (DMEM GlutaMAX™, Gibco®, 10% FCS) containing 1.6μM insulin was added and changed after 3 days. After an additional 3 days, cells were washed and maintained in regular culture medium (DMEM GlutaMAX™, Gibco®, 10% FCS) for 5 days, changing medium after 1 day and 4 days. Differentiated 3T3-L1 adipocytes were washed and control or MetR medium was added. After 48 hours, medium was aspirated, TriPure RNA Isolation Reagent (Roche Diagnostics) was added to the adipocytes, and the plate was stored at - 20°C for RNA extraction

### PVAT isolation

PVAT was carefully stripped *in vivo* from the caval vein and pooled in cold PBS. The tissue was washed by transporting into a fresh tube with PBS and thereafter weighed in separate Eppendorf tubes. After weighing, the tissue was transported into separate wells in a 24 wells-plate. Tissue was cultured under MetR or control conditions for 24 hours, medium was then transferred to an Eppendorf and stored at -80°C. Trizol was added to the tissue and the tissue was stored at -20°C for later use.

### ELISA

Mouse TNF (BD Biosciences, cat. No. 558534), IL6 (BD Biosciences, cat. No. 555240) and CCL2 (BD Biosciences, cat. No. 555260) ELISA’s were used to quantify cytokine levels in PVAT and BMDM medium according to the manufacturer’s protocol. Cytokine concentrations were corrected for the mass of PVAT of each sample.

### RNA Isolation from *Ex Vivo* and *In Vitro* Samples

*VSMCs and BMDMs* - 1mL Trizol (Invitrogen 15596026) was added to each well and pipetted up and down once before transferring to 1.5mL Eppendorf and incubating for 5 minutes at RT. *PVAT* - the tissue-samples were pooled in a 1.5mL Eppendorf, that was cooled beforehand on dry ice, and 250mL Trizol was added per Eppendorf after which the tissues were homogenized with hand-held homogenizer (Fisherbrand) and 1.5mL sterilized pestle (Axygen, PES 15-B-SI). After homogenization, the Eppendorf was centrifuged at 12.000*g* for 5 min at 4°C. Supernatant then was transferred to a new Eppendorf followed by 5 minute incubation at RT. Next, 200µL of chloroform was added, the tubes vortexed and incubated for 2 minutes on wet ice. Tubes were then centrifuged at 12.000*g* for 10 minutes at 4°C. The supernatant was collected and combined with 1µL glycogen and 500 µL isopropanol, incubated for 10 minutes at RT and centrifuged at 12.000*g* for 10 minutes at 4°C. The supernatant was discarded and 1mL 75% ethanol was added followed by centrifuged at 13.000*g*, 15 minutes at 4°C (twice). The final pellet was dried, resuspended in 20µL RNAse-free water, quantified with NanoDrop 1.000 Spectrophotometer (Thermo Scientific) and stored at -80°C. cDNA was synthesized using a High-Capacity cDNA Reverse Transcription Kit (Applied Biosystems) according to the manufacturer’s protocol. qPCR was performed on a Quant Studio 5 qPCR machine (Applied Biosystems) using Taqman gene expression assays for *Hprt1*, *Tnfα*, *Ccl2*, *Il6*, *Leptin* and *Pparα* or Sybr Green for *Tnc*, *Lox*, *Ucp1*, *Ccr2*, *Cd38*, *Gpr18* and *Arg1* according to manufacturer’s protocol.

### Primer sequences

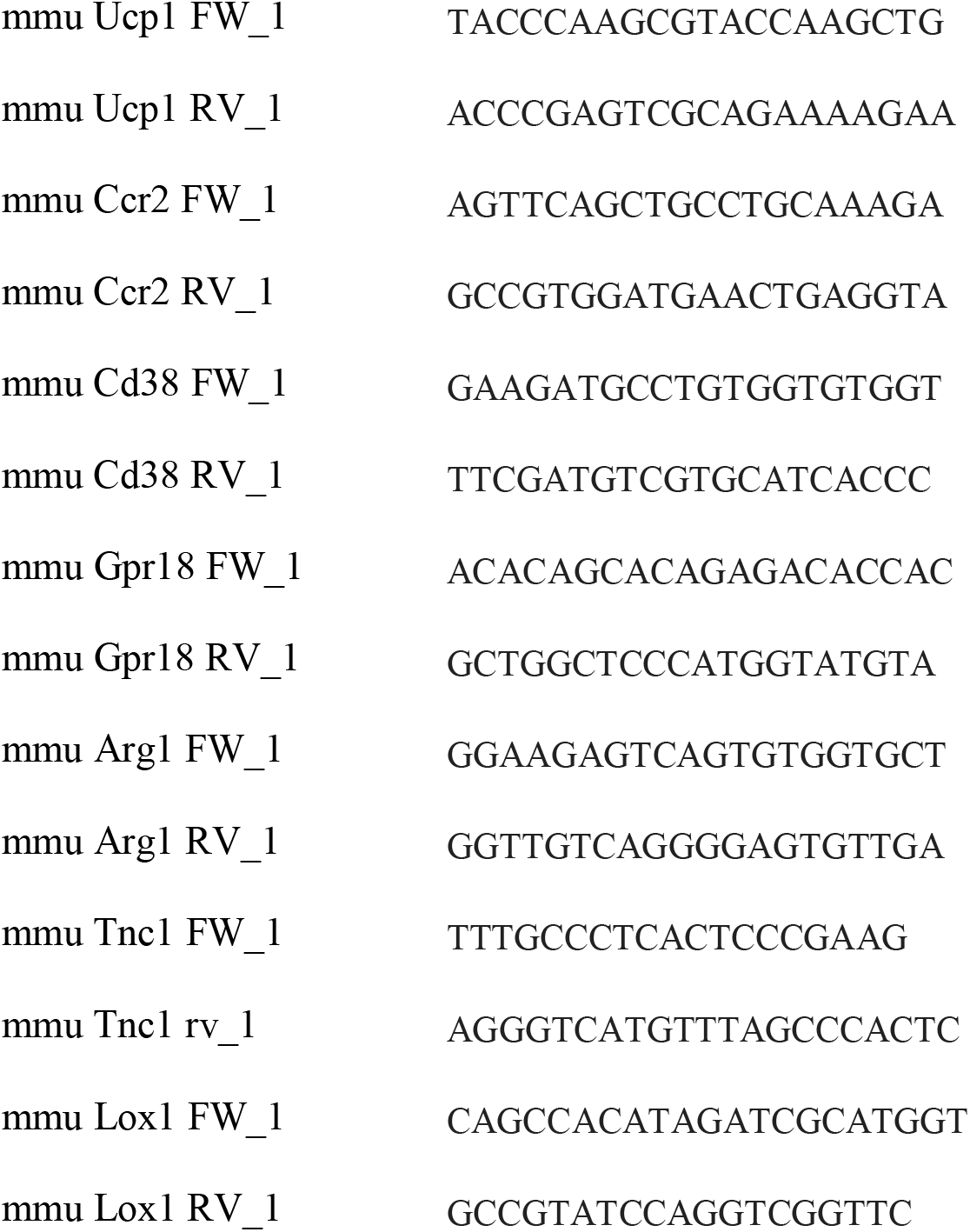

### RNA isolation from Arterial and Venous PVAT

Adipose tissue samples were collected on dry ice, snap frozen in liquid nitrogen and stored at -80°C. For RNA sequencing analysis of aorta (arterial) PVAT, the aorta PVAT of two mice on corresponding diets was pooled in one pre-cooled Eppendorf on dry ice. For caval vein and vein graft PVAT RNA sequencing, PVAT from 3 mice on corresponding diets was pooled in one pre-cooled Eppendorf on dry ice. Next, 250µL Trizol (Thermo Fisher, cat# 15596026) was added per Eppendorf and the tissue was thoroughly homogenized with a hand-held tissue homogenizer. After homogenization, samples were centrifuged at 12.000*g*, 5 minutes at 4°C. Supernatant was transfer to a fresh tube and incubated for 5 minutes at RT, then 200µL of chloroform (Sigma-Aldrich, cat#288306-1L) was added and the tube incubated for 2 minutes on wet ice. After centrifuging tubes at 12.000*g* for 10 minutes at 4°C, the aqueous layer was collected on a fresh tube on ice. 250µL iso-propanol and 1µL glycogen was added to each sample, vortexed and centrifuged (12.000*g*, 10 minutes at 4°C). Supernatant was aspirated and 75% EtOH was added before Eppendorf was vortexed and centrifuged (12.000*g*, 10 minutes at 4°C). This EtOH wash was repeated for a total of three times, then the RNA-pellet was left to dry for 20 minutes at RT. Pellet was then eluted in 20µL of RNAase free H_2_O and stored at -80°C.

### Bulk RNA Sequencing

RNA concentration and purity was measured using a Nanodrop spectrophotometer and validated with an Agilent 2100 Bio-analyzer. cDNA libraries were prepared using the Illumina TruSeq Stranded Total RNA sample preparation protocol. cDNA libraries were pooled and sequenced on an Illumina NovaSeq 6000 at a depth of approximately 20 million paired end 150bp reads per sample. Reads were aligned to the mouse GRCm38.p6 assembly using the align function and annotated using the featureCounts function from the Rsubread R package (version 2.3.7).^35^ Differential expression analysis was performed using the edgeR^36^ (3.30.3) and limma^37^ (3.44.3) R packages. ENSEMBL gene IDs were mapped to gene symbols and Entrez IDs using the mapIds function from the AnnotationDbi package (1.51.1). In total, reads were mapped to 27,179 genes. Read counts were normalized to counts per million reads and genes that did not have at least one read in at least two samples were filtered, leaving 13,667 genes. Normalization was performed using the trimmed mean of M-values method as implemented in the calcNormFactors from edgeR. Data were modeled and differential expression was determined using the limma voom pipeline to generate linear models with empirical Bayes moderation. Differential expression was determined using a Benjamini-Hochberg adjusted p-value less than 0.05. Once differentially expressed genes were determined, gene set enrichment analysis was performed using the enrichKEGG function from the clusterProfiler package (3.16.0).^38^

### Single cell Nuclear Sequencing

Nuclei from flash frozen tissues were isolated by mechanical dissociation with a glass homogenizer as previously described.^39, 40^ Homogenization was performed with a 7mL glass Dounce tissue grinder (8–10 strokes with loose pestle, 8–10 strokes with tight pestle) in homogenization buffer (250 mM sucrose, 25 mM KCl, 5 mM MgCl_2_, 10 mM Tris-HCl, 1 mM dithiothreitol (DTT), 1× protease inhibitor, 0.4 U μl^−1^ RNaseIn, 0.2 U μl^−1^ SUPERaseIn, 0.1% Triton X-100 in nuclease-free water). The nuclei suspension was filtered through a 40-μm cell strainer and centrifuged at 500*g* for 5 minutes for 4 °C. The supernatant was discarded and the pellet was resuspended in storage buffer (1× PBS, 4% bovine serum albumin (BSA), 0.2 U μl^−1^ Protector RNaseIn). Nuclei were stained with NucBlue Live ReadyProbes Reagents (ThermoFisher) and Hoechst-positive single nuclei were purified by fluorescent activated cell sorting (FACS) using a FACSAria sorter (BD Biosciences). Nuclei purity and integrity was verified by microscope. cDNA libraries were prepared by loading nuclei suspension on a Chromium Controller (10X Genomics) with a targeted nuclei recovery number of 5,000/sample followed by the Chromium Single Cell Reagent Kit v2 protocol (10X Genomics) according to the manufacturer’s protocol. Quality control of individual and pooled cDNA libraries was performed with a Bioanalyzer 4200 TapeStation System (Agilent). Pooled libraries were sequenced using 150bp paired end reads on a NextSeq 500 (Illumina) at Harvard Medical School with a minimum depth of 20,000–30,000 read pairs per nucleus. Following sequencing, Bcl files were converted to Fastq files using the Illumina bcl2fastq utility. Reads were mapped to the reference human transcriptome that was prebuilt with the 10X Genomics cellranger suite (v.3.0.1). Mapping quality was assessed using the cellranger summary statistics. Downstream analyses were performed using the Seurat R package (4.1.0).^41^

### Statistical analysis

All data are expressed as mean ± standard deviation unless indicated otherwise. Normality testing was performed employing the Shapiro-Wilk normality test. Normally distributed data was analyzed by Student’s *t-*test, one-way or two-way ANOVA. Non-normally distributed data was analyzed by Mann-Whitney test. All testing was done via Graphpad Prism (8.4.2).

## Results

### Protection from Adverse Vein Graft Remodeling via Short-term Methionine Restriction is Perivascular Adipose Tissue Dependent

We first explored whether vein graft durability could benefit from MetR preceding bypass surgery. All mice were first subjected to a 3-week 60% fat, 0% cysteine diet (Control) which contained standard levels of methionine (0.6% of diet mass, 2.6% of total protein) (**Fig. S1A**). After 3 weeks of Control diet, 1 cohort was switched to a methionine restriction (MetR) diet containing 60% fat, 0% cysteine and 0.07% diet mass of methionine, 0.3% of total protein) (**Fig. S1A**). After 1 week of MetR or continued Control diet, mice were either sacrificed and tissue harvested for caval vein /baseline studies or underwent a surgical intervention. Immediately post-operatively, all mice were switched back to the Control diet until tissue harvest (**Fig. 1A**). During the dietary intervention, mice subjected to MetR lost approximately 12% of their starting weight (**Fig. 1SB**, representative weight curve), despite hyperphagia (**Fig. S1C**), but regained weight rapidly post-operatively (**Fig. S1B**).

For the first experiment, vein graft surgery, as performed routinely^32^, included partial stripping of caval vein PVAT to better facilitate anastomosis in the recipient (**Fig. S2A**).^42^ Caval veins from donor mice were then implanted in recipients (on a corresponding diet) via an end-to-end anastomosis. At POD28, vein grafts were harvested and processed for histology (**Fig. S2B**). Morphometric analysis demonstrated that pre-operative MetR-fed mice had a significant decrease of 31% and 30.5% in I/M area and thickness ratios respectively (**Fig. S2C-D**).

We next sought to better understand the mechanism by which MetR protects against adverse vein graft remodeling. Considering the potential of PVAT to modify surgical outcome^9, 10^, together with the published effects of MetR on different adipose depots^24^, we hypothesized that protection from adverse remodeling by short-term MetR is dependent on modulation of local PVAT phenotype. For this experiment, the donor vena cava was either stripped of its PVAT (“No PVAT”) or PVAT was left intact (“PVAT”) before transplantation into a recipient on a corresponding diet (Control or MetR), thus creating a vein graft without PVAT (**Fig. 1B-C**) or completely intact PVAT (**Fig. 1D-E**). Histomorphometric analysis at POD28 (**Fig. 1F**) revealed a PVAT-dependent protection from adverse remodeling by MetR, as demonstrated by a significant decrease in I/M area ratios (53%, **Fig. 1G**), and I/M thickness ratios (53%, **Fig. 1H**) compared to Control + PVAT; while there was no difference between diet groups without PVAT. Furthermore, two-way ANOVA analysis established a significant diet-PVAT interaction in both vein graft layer thickness and area ratios (**Fig. 1I-J**). MetR + PVAT vein grafts trended towards a smaller intimal area (**Fig. S3A**) but there was no change in intimal thickness (**Fig. S3B**). At POD28, lumen area was decreased in MetR + PVAT mice (**Fig. S3C)**. The favorable morphology of MetR + PVAT vein grafts was mainly driven by an increase in media thickness, (37%, **Fig. 1I**) and area (27%, **Fig. 1J**). All data on the interaction between diet and PVAT on these histomorphometric parameters is summarized in **Table 1**.

Finally, we examined whether MetR was able to attenuate negative wall remodeling after creation of a focal stenosis in the right common carotid artery (RCCA) which induces arterial IH by altering the local hemodynamic environment (**Fig. S4A**). Control-fed mice exhibited arterial remodeling and IH after focal stenosis, while preconditioning with MetR prevented IH at POD28 (**Fig. S4B-C**).

In conclusion, MetR protects from adverse arterial and vein graft remodeling but maximal protection is dependent on the interaction between PVAT surrounding the graft and MetR.

### Short-term MetR Modulates Caval Vein Perivascular Adipose Tissue Towards an Arterial-like Phenotype

To delineate the effects of MetR on venous PVAT and to assess how MetR alters the response in PVAT to surgical injury (vein graft surgery), we performed transcriptomic analysis on caval vein PVAT (after 1 week of MetR) and on PVAT from POD1 vein grafts (**Fig. 2A**). These baseline and early time points after surgery, after MetR diet was halted, were chosen to test the hypothesis that diet/adipose interaction was present at baseline, and capable of modulating the response during early vein graft remodeling. Thoracic aorta PVAT from control-fed and MetR mice was also sequenced to investigate arterial PVAT, which could not be harvested from RCCA due to technical limitations.

**Figure 2.**
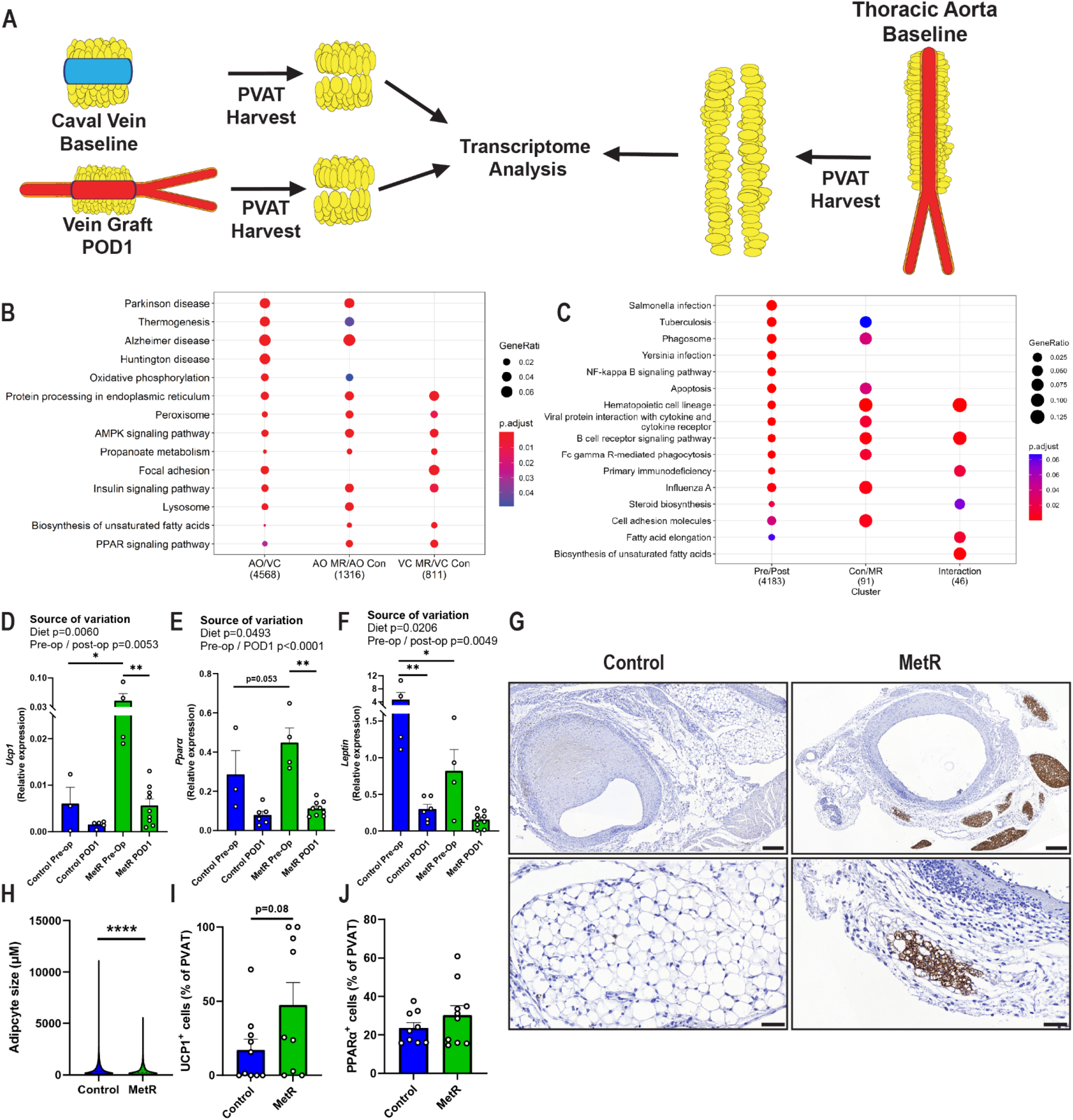
Short-term MetR Modulates Caval Vein Perivascular Adipose Tissue Towards an Arterial-like Phenotype. **A**: schematic of caval vein and vein graft POD1 PVAT; and of thoracic aorta PVAT harvest, three separate adipose tissue depots processed for transcriptome analysis simultaneously. **B**: pathway analysis of aorta versus caval vein PVAT of control-fed mice (AO/VC, first column), aorta PVAT MetR versus control-fed (AO MR/AO Con, second column) and caval vein PVAT MetR versus control-fed (VC MR/ VC Con, third column). **C:** pathway analysis of vein graft PVAT at POD1, MetR versus control-fed. **D:** *Ucp1*, **E:** *Pparα* and **F:** *Leptin* gene-transcript expression in pre-operative and POD1 PVAT, all tested via Two-way ANOVA with Bonferroni’s multiple comparisons test. **G:** Images of UCP1 stained vein grafts at POD28 in both control-fed and MetR-preconditioned mice, scale bars = 200µm (overview) or 50µm (zoom-in). **H:** individual adipocyte size in vein grafts at POD28 in both control-fed and MetR-preconditioned cohorts. **I:** Quantification of UCP1 positive cells after IHC in PVAT of vein grafts at POD28. **J:** Quantification of PPAR-α positive cells after IHC in PVAT of vein grafts at POD28. All statistical testing was done via unpaired t-test unless otherwise indicated, n=3-8/group. Graphs are presented as mean ± SEM. * P<0.05, **P<0.01, ***P<0.001, **** P<0.0001

At baseline, venous (caval vein) PVAT from control-fed mice was distinct from arterial (aorta) PVAT from control-fed mice (4568 differentially expressed genes, **Fig. S5A-B)**. Pathway analysis of these differentially expressed genes in arterial PVAT revealed a brown-adipose tissue (BAT)-like phenotype, with increased peroxisome proliferator-activated receptor (PPAR) and *thermogenesis* signatures (**Fig. 2B**). In arterial PVAT, MetR modified expression of 1316 genes (**Fig. S5C**) while further inducing PPAR and *thermogenesis* pathways (**Fig. 2B**) compared to control-fed mice.

In caval vein PVAT, MetR drove differential expression of 811 genes (**Fig. S5D**), which were enriched for PPAR signaling and fatty acid biosynthesis. (**Fig. 2B**) This transcriptomic profile, although not completely identical to arterial PVAT, does indicate increased activity of key pathways involved in browning in venous PVAT via MetR preconditioning. Taken together, these data suggest that arterial PVAT resembles BAT in terms of fatty acid catabolism and thermogenesis. Furthermore, MetR further enhances this phenotype in arterial PVAT and induces it in venous PVAT.

Analysis of PVAT at POD1 revealed vein graft surgery as a much larger modulator of gene expression than diet (**Fig. S6A)**, totaling 4183 transcripts that were differentially expressed compared to pre-operative PVAT (**Fig. S6B)**, which mainly involved pathways that regulate innate and adaptive immune response (**Fig. 2C**). When comparing POD1 PVAT, diet differentially modulated 91 transcripts (**Fig. S7B**), which further enhanced regulation of immune function pathways in this adipose depot (**Fig. 2C**). Interestingly, we found several pathways that displayed a strong interaction between diet and surgery, with regulation confined to a solitary diet group following surgery (**Fig. 2C**). These pathways were either involved in immune function (hematopoietic cell lineage, B-cell receptor signaling, primary immunodeficiency) or lipid- and steroid hormone biosynthesis (**Fig. 2C**), suggestive of a diet-dependent immune response to the surgical stress. In line with these findings, we observed a dampening of Toll-like receptor 4 (*Tlr4*) transcript expression (**Fig. S6C**) together with a robust decrease in expression of its endogenous ligand^43^ tenascin-C (**Fig. S6D**) in the MetR group. Lysyl-oxidase^44^ (*Lox*), an enzyme involved in atherogenesis and restenosis, was also strongly downregulated in MetR PVAT compared to control-fed at POD1 (**Fig. S6E**).

Since we observed general PPAR signaling as an effect of MetR in venous PVAT pre-surgery, we also investigated specific genes involved in browning (i.e., *Ucp1*, *Pparα*) in PVAT. *Ucp1* expression was increased by MetR in pre-op PVAT and also significantly altered as a function of vein graft surgery (**Fig. 2D**). Although we did not detect significant differential expression of *Pparα* as a function of diet at pre-op PVAT, there was a significant main effect of diet on expression across surgery conditions (**Fig. 2E**). Expression of *Leptin* was significantly decreased by MetR in pre-op PVAT. In addition, surgery as well as diet significantly lowered *Leptin* expression. (**Fig. 2F**). In POD28 vein grafts, both an increase in UCP1 protein levels (77% increase, **Fig. 2G-I**) and a decrease in adipocyte size^45^ (**Fig. 2H**) were suggestive of a temporal browning phenotype (**Fig. 2F-H**), while PPAR-α expression differences were absent at this late-stage remodeling timepoint (**Fig 2J**).

All in all, these data suggest a MetR-dependent and transient activation of thermogenesis and PPAR-α signaling in venous PVAT at both baseline and during early vein graft remodeling, resembling an arterial PVAT phenotype. Moreover, in response to the vein graft surgery, PVAT innate and adaptive immune systems were strongly activated, and diet appeared to modulate or dampen this response.

### Short-Term MetR Induces Browning in Venous PVAT via Adipocyte Specific PPAR-*α* Activation and Dampens the Post-operative Pro-Inflammatory Macrophage Response

To determine which specific cell type in MetR-preconditioned PVAT is involved in thermogenic activation and immune response modulation, and to better understand which cell-specific pathways are involved in protection from vein graft remodeling, we performed single-nucleus RNA sequencing on control-fed and MetR preconditioned PVAT from caval veins at baseline and from vein grafts at POD1. **Fig. S7A-C** demonstrates technical validity of the sequencing, with low per-sample and inter-sample variability. Both Control and MetR PVAT had a distinct UMAP cell clustering signature, both at the pre-operative and POD1 time-point (**Fig. 3A-B**). We identified robust marker genes for each cluster with limited redundancy between clusters (**Fig. 3C**). Cell type identity for each cluster was assigned using canonical marker genes (**Fig. S8A**) which was validated using an unbiased comparison to reference datasets (**Fig. S8D**). **Fig. S8E** compares the proportion of each cell cycle state across clusters.

**Figure 3.**
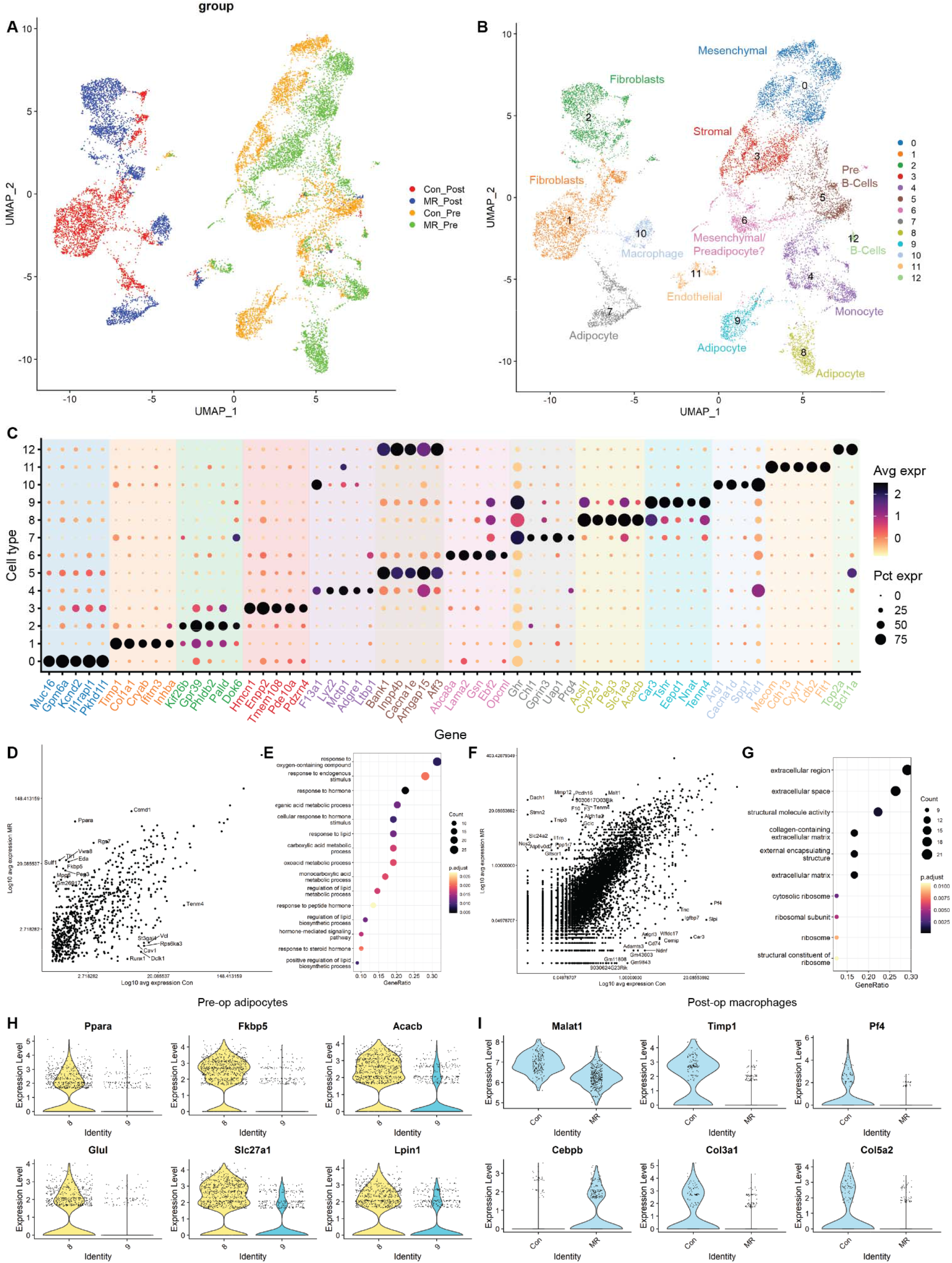
Pre-operative MetR induces browning in PVAT adipocytes via PPAR-α activation and dampens the post-operative pro-inflammatory macrophage response. Single cell nuclear sequencing was performed on IVC and vein graft POD1 PVAT in control-fed and MetR mice, with n=2 samples processed per group. **A:** UMAP project of cells colored by diet/surgery group. N=2 samples per group (Control/MetR (Con/MR) and pre-operatively/postoperatively (Pre/Post) . **B:** UMAP projection of cells colored by cluster with cell type description. N=2 samples per group **C:** Top 5 most characteristic genes for each cluster. **D:** Expression of genes in pre-op adipocytes from control (cluster 9) and MetR (cluster 6). **E:** Pathway analysis of top 100 differentially expressed genes in pre-op adipocytes between control and MetR groups. **F:** gene expression in post-op macrophages (10) in control and MetR groups. **G:** pathway analysis of top 100 differentially expressed genes in post-op macrophages between control vs MetR. **H:** Violin plots of top differentially expressed genes in pre-op adipocytes. **I:** Violin plots of top differentially expressed genes in post-op macrophages.

Pre-conditioning with MetR resulted in pronounced changes in transcriptional profiles in several PVAT cell types (**Fig. 3A-B**), including mesenchymal (cluster 0) and stromal cells (cluster 3). The greatest pre-operative signature change after MetR however, was seen in adipocytes to the extent that they were classified into distinct clusters (cluster 8-9). In pre-operative PVAT adipocytes (**Fig. 3D**), several pathways involved in energy- and fatty-acid metabolism were highly differentially regulated in the MetR group (**Fig. 3E**). Notably, there was marked increased expression of *Pparα* and several related gene transcripts and this effect was confined to MetR preconditioned pre-operative PVAT adipocytes (**Fig. 3H**).

At the POD1 time-point, several clusters showed distinct transcriptional signatures induced by diet (**Fig. 3A-B**), including fibroblasts (cluster 1-2), adipocytes (cluster 7) and macrophages (cluster 10). In post-operative macrophages (**Fig. 3F**), preconditioning with MetR resulted in marked upregulation of pathways involved in the formation of extra-cellular matrix proteins and structures (**Fig. 3G**). Individual transcripts that were shown to be highly regulated by MetR in post-operative macrophages were mainly associated with an anti-inflammatory macrophage phenotype, such as *Malat1*^46^ and *Cebpd*^47^ (**Fig. 3I**). Interestingly, PPAR-α regulation was not present anymore in MetR POD1 PVAT adipocytes, but rather upregulation of several pathways involved in extracellular matrix remodeling and protein synthesis, as revealed by the POD1 adipocyte pathway analysis (**Fig. S8E**).

Collectively, these data are suggestive of a central role for PPAR-α to induce the “arterial-like” phenotype seen in MetR preconditioned venous PVAT, and that this highly transient signature of PPAR-α activation in PVAT adipocytes is already absent at POD1. At POD1, PVAT of vein grafts that received pre-operative MetR had a marked anti-inflammatory phenotype specifically in macrophages, via *Malat1* and *Cebpb* transcript regulation (**Fig 3F-G, I**).

### MetR Activates PPAR-*α* Signaling in Adipocytes and Dampens the Inflammatory Response of Macrophages and PVAT *In vitro*

To assess the direct effect of MetR on secretion of pro-inflammatory cytokines by PVAT, we incubated venous PVAT in either control or MetR culture medium and found decreased secretion of IL-6 (**Fig. 4A**), CCL2 (**Fig. 4B**) en TNF-α (**Fig. 4C**) in the supernatant of MetR preconditioned PVAT, confirming the anti-inflammatory phenotype of PVAT in response to MetR *in vitro*.

**Figure 4.**
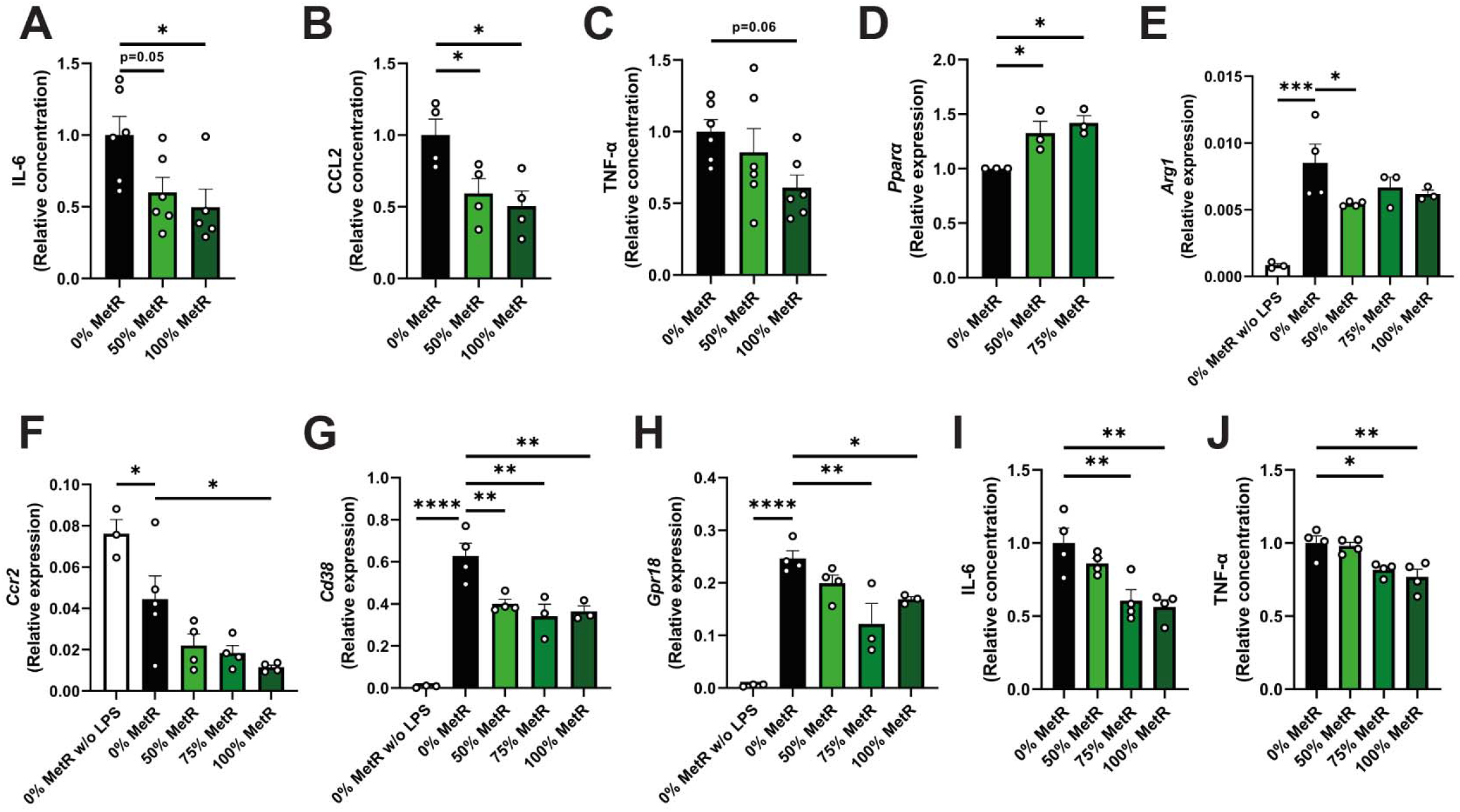
MetR Activates PPAR-α Signaling in adipocytes and Dampens the Inflammatory Response of Macrophages and PVAT In vitro. **A-C:** venous PVAT was harvested from C57BL/6 mice and incubated for 24-hours in either complete-, 50%- or 100% MetR medium, supernatant was then interrogated via ELISA. **A:** IL-6 levels in supernatant of PVAT. **B:** CCL2 levels in supernatant of PVAT. **C:** TNF-α levels in supernatant of PVAT. **D:** qPCR for *Pparα* in adipocyte cell line (3T3-L1 cells) after 24-hour incubation with MetR. **E-H**: bone-marrow derived macrophages were harvested from C57BL/6 mice and stimulated with LPS to create a pro-inflammatory state, followed by 24-hour MetR. **E:** qPCR for *Arg1* in pro-inflammatory macrophages under control and MetR conditions. **F:** qPCR for *Ccr2* in pro-inflammatory macrophages under control and MetR conditions. **G:** qPCR for *Cd38* in pro-inflammatory macrophages under control and MetR conditions. **H:** qPCR for *Gpr18* in pro-inflammatory macrophages under control and MetR conditions. **I-J:** supernatant from previous experiment tested for cytokine production via ELISA. **I:** IL-6 production. **J:** TNF-α production. All statistical testing was done via one-way ANOVA with Dunnet’s multiple comparisons test, unless otherwise indicated, n=3-6/group. Graphs are presented as mean ± SEM. * P<0.05, ** P<0.01, *** P<0.001, **** P<0.0001.

Next, we sought to test PPAR-α activation specifically in adipocytes using a 3T3-L1 adipocyte cell line. Incubation of these adipocytes with MetR-medium showed a significant induction of PPAR-α (**Fig. 4D**).

Thirdly, we cultured bone-marrow derived macrophages from C57BL/6 mice and stimulated them with LPS to induce a pro-inflammatory phenotype. Next, we subjected them to various concentrations of MetR to interrogate its effect on pro-inflammatory macrophages. Interestingly, *Arg1* (**Fig 4E**, anti-inflammatory macrophage marker^48^) was still modestly downregulated, and we did not find any difference in *Tnfα* and *Il6* transcripts (**Fig. S9A-B**). There was however a strong decrease of *Ccr2-* (**Fig. 4F**), *Cd38-* (**Fig. 4G**, pro-inflammatory macrophage marker^49^) and *Gpr18-*transcripts (**Fig. 4H**, pro-inflammatory macrophage marker^50^) compared to control-medium, suggesting a diminished pro-inflammatory response after LPS stimulus. Finally, we tested the supernatant of these macrophages after LPS and MetR incubation and found diminished production of the pro-inflammatory cytokines IL-6 (**Fig. 4I**) and TNF-α (**Fig. 4J**), but not CCL2 (**Fig. S9C**)

To summarize, we were able to replicate our *in vivo* findings by showing dampening of PVAT pro-inflammatory cytokine production by MetR *ex vivo,* and direct *in vitro* PPAR-α activation in MetR-treated adipocytes. Secondly, interrogation of pro-inflammatory macrophages in a MetR environment recapitulated the macrophage shift towards an anti-inflammatory phenotype at the transcriptional and cytokine secretion levels.

### Increased Polarization of Macrophages Towards an Anti-inflammatory Phenotype is Driven by the MetR-PVAT Interaction in Vivo

To better understand whether any of these pre-surgery and early remodeling findings in macrophages and adipocytes could be linked to a late-stage favorable remodeling phenotype, we next interrogated vein grafts at POD28. In the MetR + PVAT group, reduced pro-/anti-inflammatory macrophage ratios were observed both in the graft wall (**Fig. 5A, C**) and PVAT (**Fig. 5B,C**). This favorable ratio present in both vein graft (VG) layers was driven by a significant interaction between MetR and the presence of PVAT (**Fig. 5A, Table 2**). The observed differences in ratios could be explained by increased polarization towards the anti-inflammatory phenotype, both in the intimal and media layers (**Fig. 5D**) and in MetR PVAT (**Fig. S10A**), while there was a limited decrease in pro-inflammatory macrophages in the vein graft wall (**Fig. 5E**) and PVAT (**Fig. S10B)**. Preconditioning with MetR yielded no difference in total M_□_ macrophages in the vein graft wall (**Fig. S10C**), nor in surrounding PVAT (**Fig. S10D**). This increase in intimal anti-inflammatory macrophages together with an apparent small decrease in pro-inflammatory macrophages in the MetR + PVAT cohort could also be linked to the underlying interplay between diet and PVAT (**Fig. 5D-E, Table 2**).

**Figure 5.**
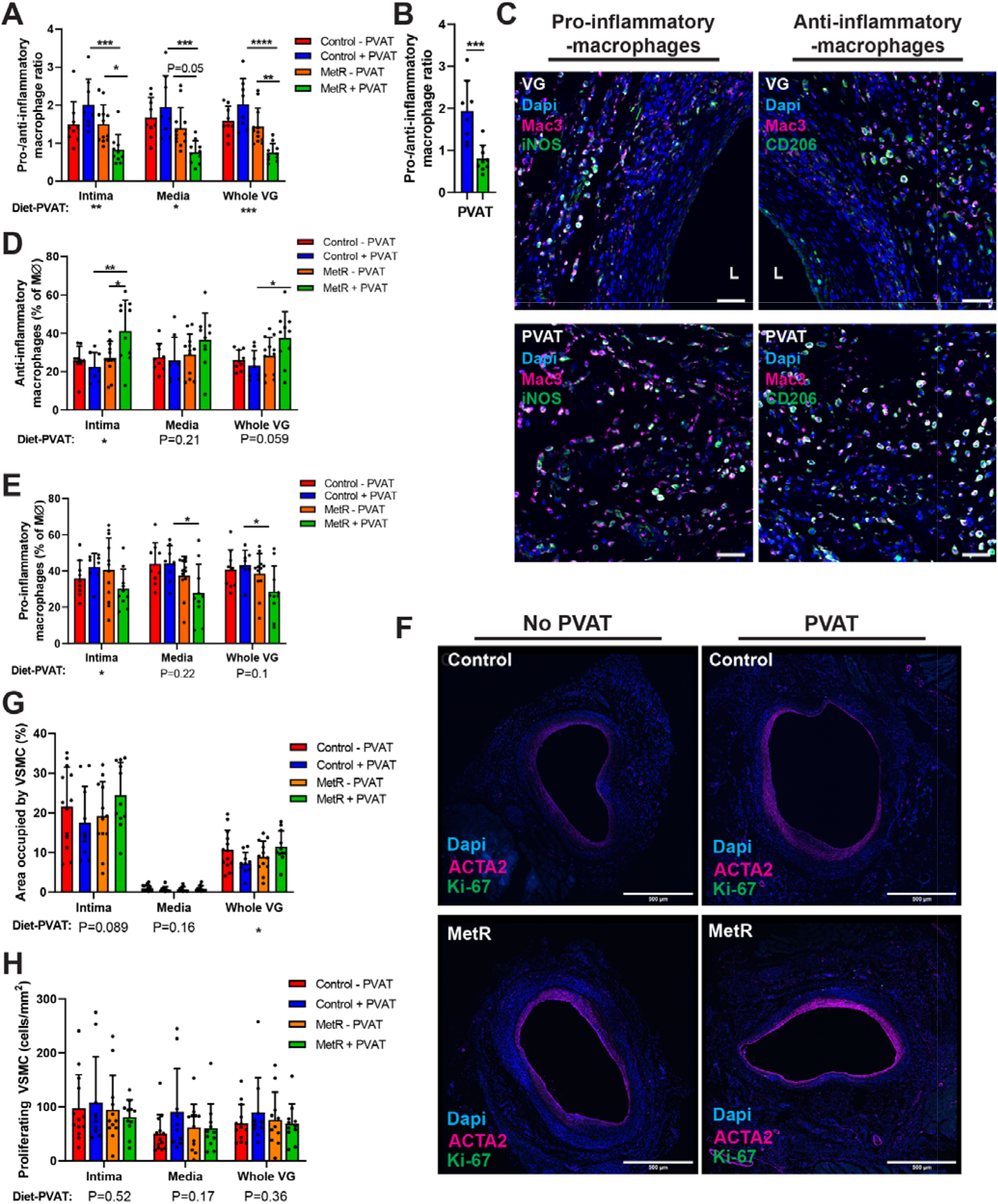
Favorable Vein Graft Wall Composition Seen in MetR Mice is Also Perivascular Adipose Tissue Dependent. **A:** Pro-/anti-inflammatory macrophage ratio per vein graft layer. **B:** Pro-/anti-inflammatory macrophage ratio in PVAT of MetR and control-fed mice. **C:** immunohistochemical staining for pro- and anti-inflammatory macrophages, L indicates lumen. Scale bars = 500µm. **D**: Anti-inflammatory macrophages as a percentage of total MΦ per vein graft layer. **E**: Pro-inflammatory macrophages as a percentage of total MΦ per vein graft layer. **F**: immunohistochemical staining for ACTA2 and Ki-67. Scale bars = 500µm **G:** area occupied by VSMC per vein graft layer. **H:** proliferating VSMC (ACTA2 + Ki-67 double positive cells) per mm^2^ per vein graft layer. All statistical testing was done via two-way ANOVA with Tukey’s multiple comparisons test, unless otherwise indicated, n=10-13/group. * P<0.05, ** P<0.01, *** P<0.001, **** P<0.0001.

**Figure 6.**
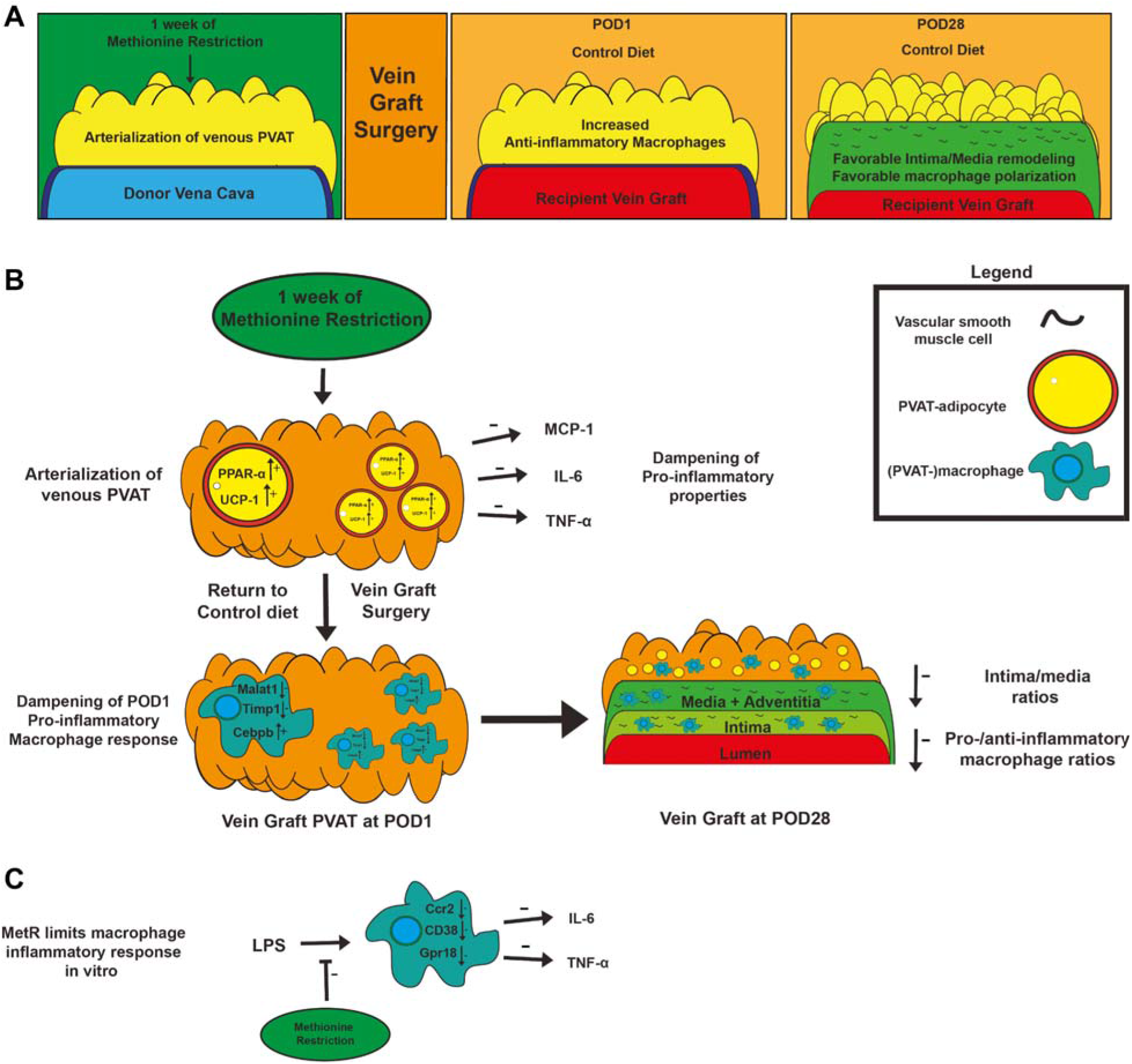
Methionine Restriction Protects from Adverse Vein Graft Remodeling via Pro-inflammatory Macrophage Modulation and PPAR-α Induced Browning in Perivascular Adipose Tissue. **A:** General overview of proposed mechanism of MetR-PVAT interaction dependent protection from VGD. **B:** In vivo/*ex vivo* mechanisms of MetR-PVAT interactions at pre-op, POD1 and POD28 timepoints. **C:** Effects of MetR on macrophages in vitro

Also, at POD28, percentage of intimal area and whole VG occupied by VSMC was dependent on the interaction between diet-PVAT (**Fig. 5G-F, Table 3**), but no significant difference in directionality could be detected between groups. The absolute number of VSMC per mm^2^ present was comparably dependent on a combination of diet and PVAT, but this effect was not strong enough to display significant inter-group differences (**Fig. S10E, Table 3**). There was no detectable change in proliferating VSMC at POD28 (**Fig. 5G, Table 3**), nor did extra-cellular matrix analysis reveal a change in collagen deposition (**Fig. S10F, Table 3**).

Taken together these data reveal that short-term MetR transformed venous PVAT towards a more arterial-like state by induction of browning and specifically upregulated PPAR-α in PVAT-adipocytes. This transition to a more arterial-like transcriptional profile was accompanied by an early anti-inflammatory response in PVAT-macrophages and resulted in favorable late-stage vein graft remodeling phenotype with favorable anti/pro-inflammatory macrophages ratios as the main driver in protection from graft failure.

## Discussion

Here we tested the potential of short-term restriction of the sulfur amino-acids methionine and cysteine (MetR) in preclinical models of cardiovascular surgery interventions. Excitingly, just one week of the MetR intervention, which constitutes an isocaloric diet with adequate levels of all macronutrients, protected from arterial IH in a focal stenosis model, and protected from adverse remodeling after vein graft surgery. Specifically in bypass surgery, we show that protection from MetR was entirely dependent on the presence of PVAT. Control-fed mice who received a caval vein with intact PVAT had exacerbated vein graft disease (VGD) at POD28, pointing towards a diet-induced reversal of PVAT phenotype in MetR mice compared to control diet. There was no significant histomorphometric difference detectable between diet groups when caval veins were stripped of PVAT before anastomosis creation, indicating a PVAT-dependent protection from negative remodeling yielded by MetR.

In this study, we performed a side-by-side comparison of arterial and venous PVAT via whole-transcriptome sequencing, whilst also examining the effects of MetR and surgical stress on these PVAT phenotypes. Our study revealed a distinctly separate transcriptomic profile between vein and artery PVAT, with venous PVAT resembling white/beige adipose tissue.^51^ Arterial PVAT resembles BAT, as seen by the presence of several genes involved in thermogenesis and in accordance with previous studies.^52^ Interestingly, short-term MetR further increased browning in arterial PVAT, while it induced this phenotype in venous PVAT, which was further confirmed with *in vivo* and *in vitro* experiments. And while it is known that long-term (weeks-months) MetR induces browning in inguinal beige adipose tissue^25, 53, 54^, here we were able to induce this phenotype in venous PVAT after just 1 week of MetR. Transcriptomic changes in response to surgery however were much larger than in response to diet at the POD1 timepoint, which makes sense considering the severe changes in environment and stressors that veins undergo when switched to an arterial circulation.^4^ We noted strong activation of several pathways involved in the immune response, which confirms previous findings in vein grafts regarding the early remodeling stages and involvement of immune-cells.^55–57^

Next to whole-transcriptome sequencing, we also performed PVAT analysis on a single-cell whole-transcriptome level. Our analysis revealed that browning of PVAT coincided with upregulation of PPAR-α in adipocytes upon MetR, which corroborates evidence from previous studies on long-term MetR in inguinal adipose tissue.^25, 58^ This signature was highly transient and not present in POD1 PVAT-adipocytes, therefore likely a direct result of MetR. Whether this concerns a cell-autonomous effect of MetR on PVAT-adipocytes or signaling via other cell-types or organs is not fully determined. It is likely that MetR activates PPAR-α locally and directly in PVAT-adipocytes since we were able to replicate this *in vitro* in adipocytes under MetR conditions. A recent study by Decano et al.^59^ revealed a novel and decisive role for PPAR-α in VGD, demonstrating prevention of graft failure by PPAR-α activation specifically in macrophages, which muted their pro-inflammatory capabilities. Via gain- and loss- of function vein graft experiments, they show that PPAR-α activation in macrophages muted their pro-inflammatory capabilities via increased scavenging of reactive-oxygen-species and inhibition of nitric oxide synthase. Additionally, loss of PPAR-α was recently found to be associated with negative vein graft remodeling by differentiation of PVAT-derived mesenchymal stem cells into SMCs which contribute to IH, pointing towards a pivotal role of PPAR-α signaling in various cell types.^60^ Our results suggest that PPAR-α may also play a protective role in PVAT adipocytes, strengthening the evidence for an overall beneficial role of PPAR-α signaling within the vein graft microenvironment.

Next to local activation of adipocytes, it is possible that PVAT browning is also induced via a signaling pathway in response to MetR. Previous studies show that under conditions of low methionine, the liver produces fibroblast growth factor -21 (FGF-21)^61^, a hormone directly responsible for regulation of thermogenesis in both white adipose tissue and BAT depots.^62^ In the current study we subjected both donor and recipient to MetR before vein graft surgery, leaving open the possibility that browning is induced both via a paracrine effect of MetR on adipocytes in donor-PVAT, and increased hepatic FGF-21 production in the recipient.^62^

Our MetR cohort lost approximately 10% of their body weight during the course of the diet, while they significantly increased their calorie-intake. This finding is consistent with previous studies.^30, 31^ Previous work has established that the weight loss observed in MetR-fed mice is due to increased energy expenditure rather than a decrease in calorie intake.^63^ While we cannot fully rule out that part of the diet-induced benefits is due to a state of relative calorie deficit; even in the presence of hyperphagia there is an increased energy expenditure throughout the course of the MetR-diet. Of importance, when compared side-to-side MetR and calorie restriction invoke unique metabolic and inflammatory transcriptomic profiles in liver^64^, suggestive of distinct pathway activation between calorie restriction and MetR.

Single cell analysis of vein graft PVAT at POD1 revealed a marked shift in macrophages towards an anti-inflammatory phenotype after MetR, deducted from *Malat1* and *Timp1* downregulation *in vivo*. Re-iterating these findings *in vitro*, MetR conditions shifted cultured macrophages towards an anti-inflammatory phenotype, with the net result of decreased production of the pro-inflammatory cytokines IL-6 and TNF-α. In addition, analysis of vein grafts at POD28 revealed a marked shift in macrophage polarization towards the anti-inflammatory phenotype in the MetR+PVAT group, and this was dependent on the diet-PVAT interaction.

In adverse vein graft remodeling, macrophages function as one of the main effector cells in the cascade ultimately leading to graft occlusion^4^, and the presence of anti-inflammatory macrophages in grafts is associated with favorable remodeling signatures.^32^ When methionine is in abundance, macrophages progress towards a pro-inflammatory phenotype^65^, but here we show a robust reversal when methionine is restricted. Increased polarization of macrophages towards an anti-inflammatory phenotype is associated with low levels of TIMP-1^66^, while *MALAT1* is a long-coding RNA^46^ that recently has been inversely correlated with in-stent stenosis after coronary revascularisation.^67^

It is thought that *MALAT1* functions as a negative feedback sensor for the activation of the TLR4 pathway in macrophages in response to endogenous or exogenous ligands.^68^ Whether MetR directly downregulates *Malat1* or that this is merely a result of decreased activation of pro-inflammatory pathways (such as TLR4) within macrophages preconditioned with MetR is unknown. Interestingly, our whole-transcriptome data reveal marked downregulation of *Tlr4* and the TLR4 ligand Tenascin-C, suggesting the latter. In the context of adverse vein graft remodeling, deficiency of TLR4 is directly linked to a decrease in adverse graft wall remodeling.^32, 69^ Finally, whether this macrophage shift towards an anti-inflammatory state is cell autonomous in response to MetR, or whether this is driven by paracrine signaling within the vein graft microenvironment (perhaps via PVAT-adipocytes) is unknown.

In (coronary) bypass surgery, PVAT has been previously studied with regard to the “no touch” technique. In this approach, the vein graft is transplanted with its PVAT intact to potentially improve graft patency.^6, 70, 71^ Recently, several trials in humans have been conducted but with conflicting results. A 2019 trial by Deb et al. was not able to find any differences in patency rates between groups^72^, while a 2021 trial by Tian et al. did report over 40% reduction in vein graft occlusion in the no-touch group.^73^ In a study performed by Deb et al., average BMI in both groups was 28-29, while Tian et al. noted significantly diminished patency rates in high-BMI (>25) cohorts after sub-group analysis.

High-fat feeding and subsequent obesity is known to have detrimental effects on PVAT-adipocytes and phenotype, diminishing vascular (wall) function and health.^7, 74–76^ Specifically in cardiovascular surgery models, high-fat feeding exacerbates intimal hyperplasia after vascular injury via pro-inflammatory changes in PVAT.^8, 9, 77^ Extrapolating these findings to the aforementioned clinical trials, high BMI could be a proxy for the presence of an unfavorable PVAT-phenotype with regards to vein graft patency, perhaps part of the explanation for these conflicting results. In our study, we also noticed detrimental effects on PVAT-phenotype and subsequential vein graft remodeling after (prolonged) Control-diet high-fat feeding, but this was completely reversed by short-term MetR compared even to MetR without PVAT.

In summary, here we demonstrate that short-term MetR induces browning of PVAT and holds promise to improve post-interventional vascular remodeling by mitigating the detrimental effects of dysfunctional PVAT. This low-cost, simple dietary intervention, which is feasible and safe in humans,^28^ seems therefore suitable for evaluation in cardiovascular surgery clinical settings.

## Abbreviations

BAT: Brown Adipose Tissue
BMDM: Bone Marrow-Derived Macrophages
CCL2: C-C motif ligand 2
DR: Dietary Restriction
FGF21: Fibroblast Growth Factor 21
HFD: High Fat Diet
I / M: Intima / Media
IH: Intimal Hyperplasia
IL: Interleukin
MetR: Methionine Restriction
POD: Post-op Day
PPAR: Peroxisome proliferator-activated receptor
PVAT: Perivascular Adipose Tissue
RCCA: Right Common Carotid Artery
TLR4: Toll-like receptor 4
TNF-α: Tissue necrosis factor α
UCP1: uncoupling protein 1
VSMC: Vascular Smooth Muscle Cell

## Acknowledgements

P.K. provided funding, conceived of experimental designs, performed experiments and wrote the manuscript. T.S. performed *in vitro*/*in vivo* experiments and data analysis. M.T. performed surgeries, processed vein grafts and analyzed histology. P.H.Q. advised on analysis of vein grafts. J.W.J. assisted with *ex vivo* and *in vitro* experiments. J.G. and J.S. performed and advised on single-cell nuclear sequencing experiments. M.R.M. and S.J.M. assisted with animal care and *in vivo* studies, M.R.M performed data analysis in bulk and nuclear RNA sequencing. S.K. advised on analysis of *in vitro* experiments, N.K. performed *in vitro* experiments. M.A. advised on data interpretation. M.R.V., J.R.M. and C.K.O. provided funding, conceived of experimental designs and supervised the project. All authors reviewed and approved the final version of the submitted manuscript.

