## Supplementary table for "Short-term Pre-operative Methionine Restriction Induces Browning of Perivascular Adipose Tissue and Improves Vein Graft Remodeling in Mice"

**Supplemental materials**

***Supplemental figures S1-S10***

***Tables 1 – 3***

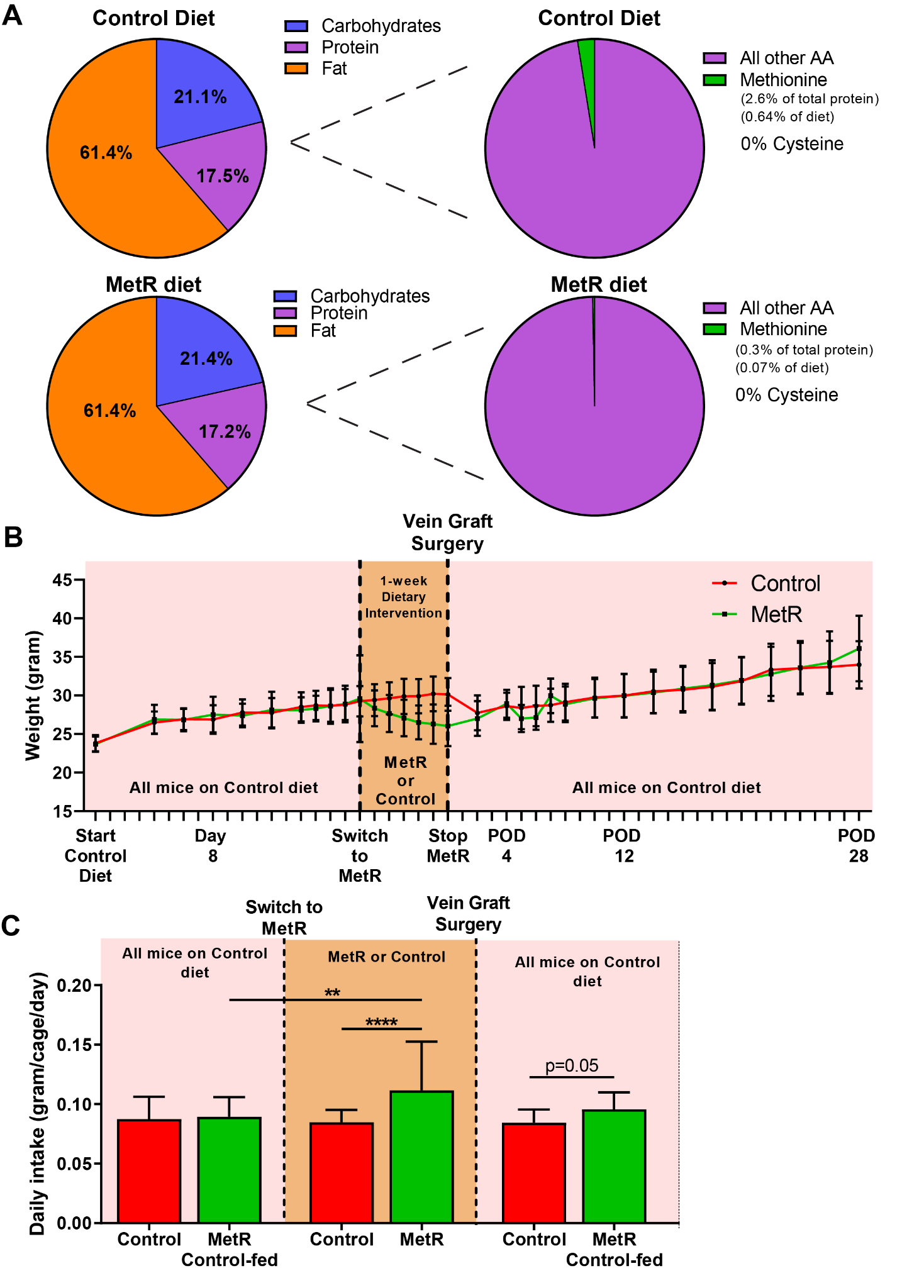

**Supplemental Figure 1.** *Methionine restriction diet composition and metabolic response.*

**A:** Macronutrient composition of control and methionine restricted diet. Both diets contain 0% cysteine and 60% fat. **B:** representative graph of weight (in gram) that was lost/gained during the study. **C**: representative bar graph of daily food-intake during study in both groups. ****** P<0.01, ******** P<0.0001. All statistical testing was done via two-way ANOVA with Tukey's multiple comparisons test, unless otherwise indicated.

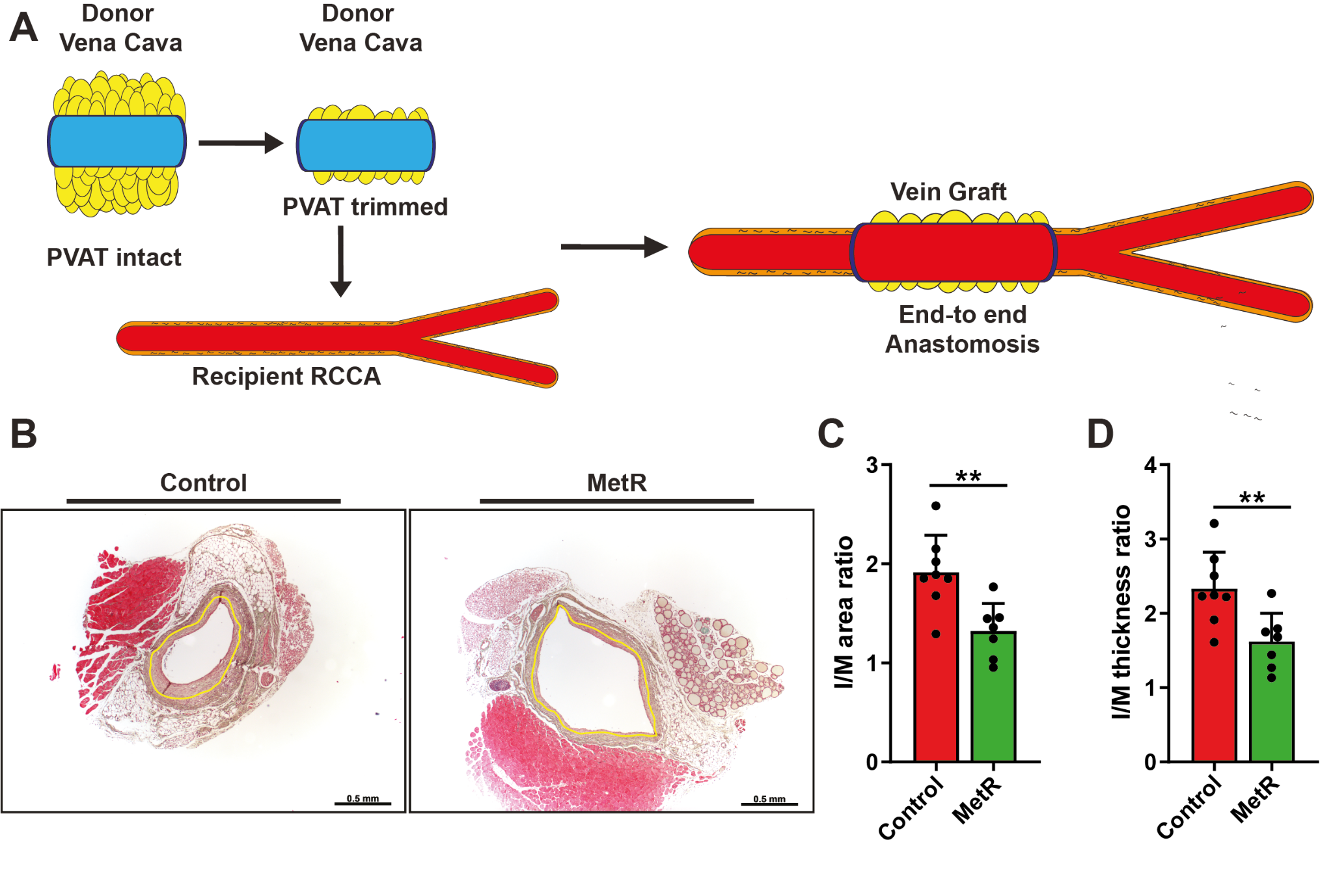
**Supplemental Figure 2.** *MetR in vein graft surgery.*

**A:** vein graft surgery procedure, with partial trimming of PVAT from donor vena ceva. **B**: vein grafts at POD28 after Masson-trichrome staining. Yellow lining indicates internal elastic lamina. Scale bars = 0.5mm. **C**: I/M area ratio with Student’s t-test, n=7-8/group. **D**: I/M thickness ratio with Student’s t-test, n=7-8/group. ****** P<0.01

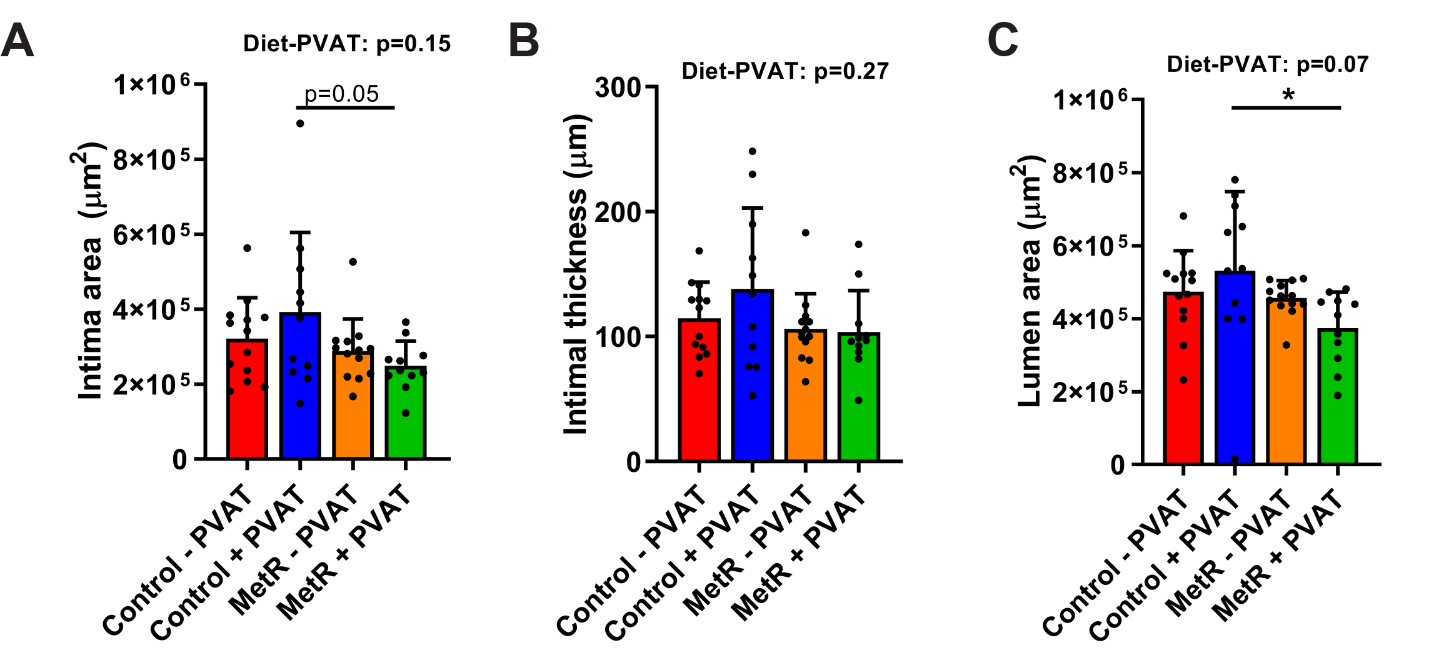
**Supplemental Figure 3.** *Intimal area/thickness and lumen diameters at POD28*.

**A-C:** Histomorphometric analysis of vein grafts at POD28. **A**: Intimal area. **B**: Intimal thickness. **C:** Lumen area. All statistical testing was done via two-way ANOVA with Tukey’s multiple comparison’s test unless otherwise indicated, n=11-13/group. ***** P<0.05

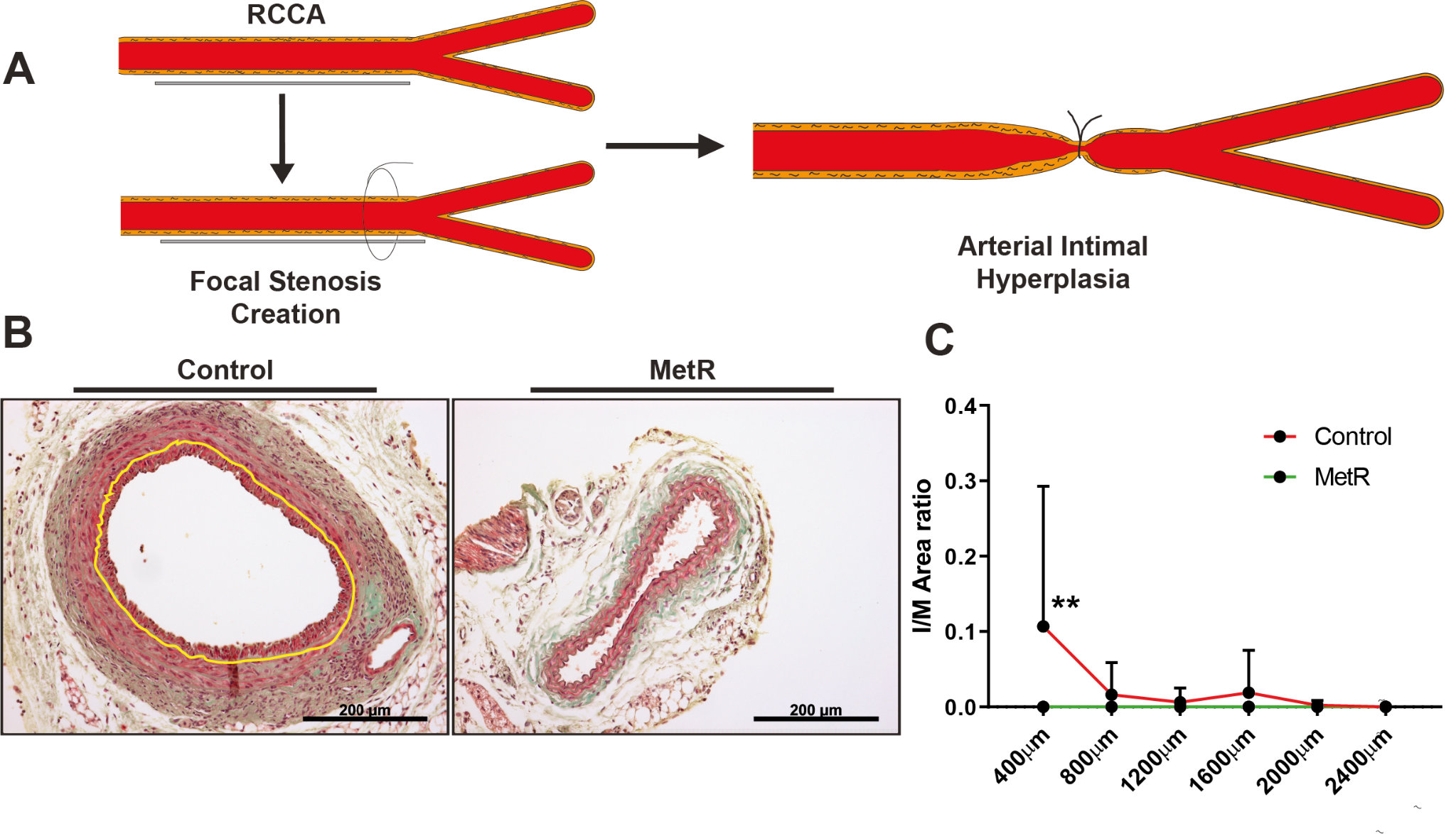

**Supplemental Figure 4.** *MetR in arterial intimal hyperplasia.*

**A:** focal stenosis creation. **B**: RCCA at POD28 after Masson-trichrome staining. Yellow line indicates internal elastic lamina lining. Scale bars = 200µm. **C:** I/M area ratio at POD28, via Two-way ANOVA with Sidak’s multiple comparisons test, n=8-9/group. ** P<0.01

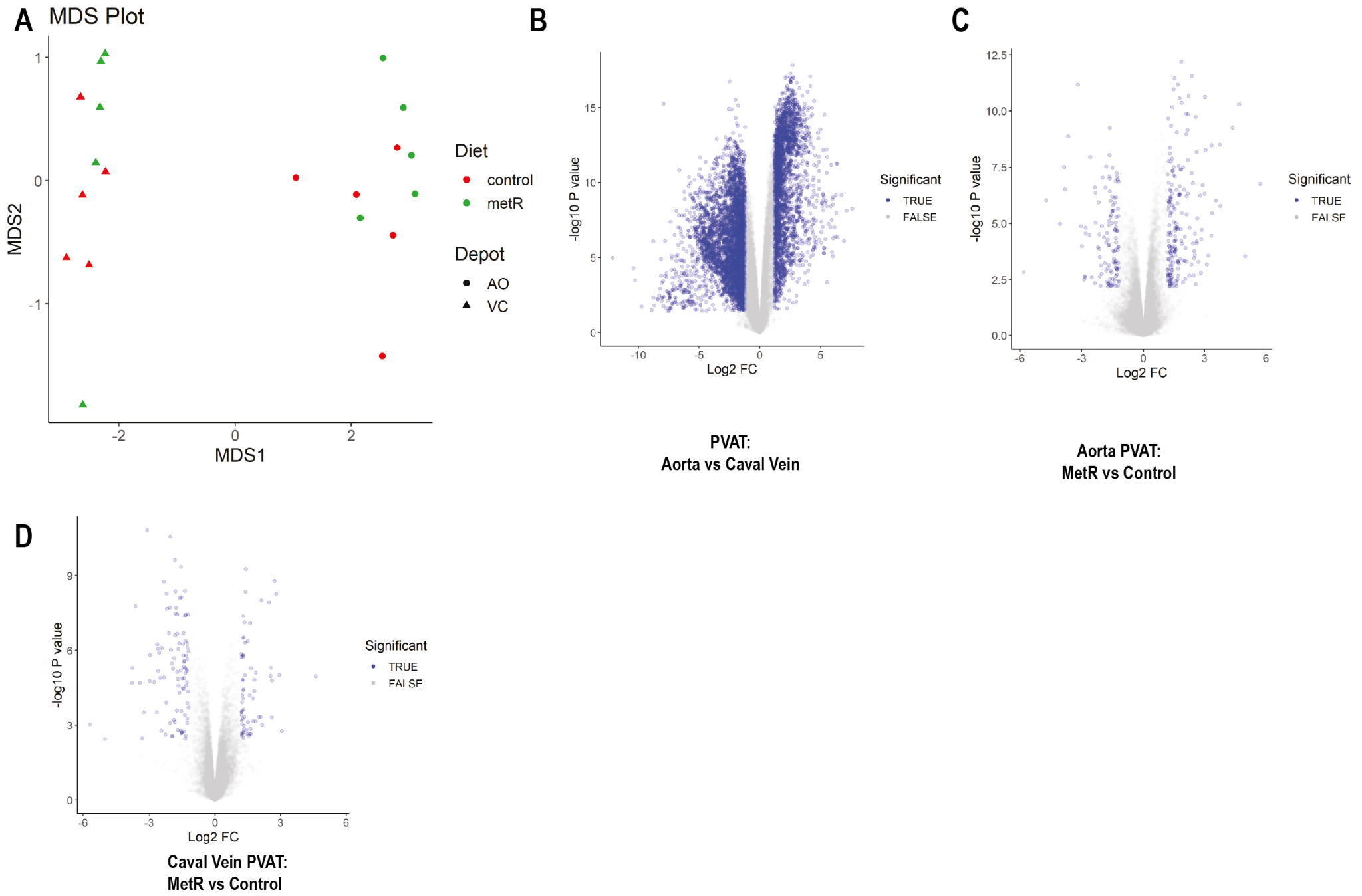

**Supplemental Figure 5.** *Baseline transcriptome analysis in arterial and venous PVAT.*
**A**: principal component analysis of aorta and caval vein PVAT of control-fed and MetR mice. **B:** fold change in transcript expression in control-fed aorta PVAT versus control-fed caval vein PVAT. **C**: fold change in transcript expression in aorta PVAT of MetR versus control-fed mice. **D**: fold change in transcript expression in caval vein PVAT of MetR versus control-fed mice.

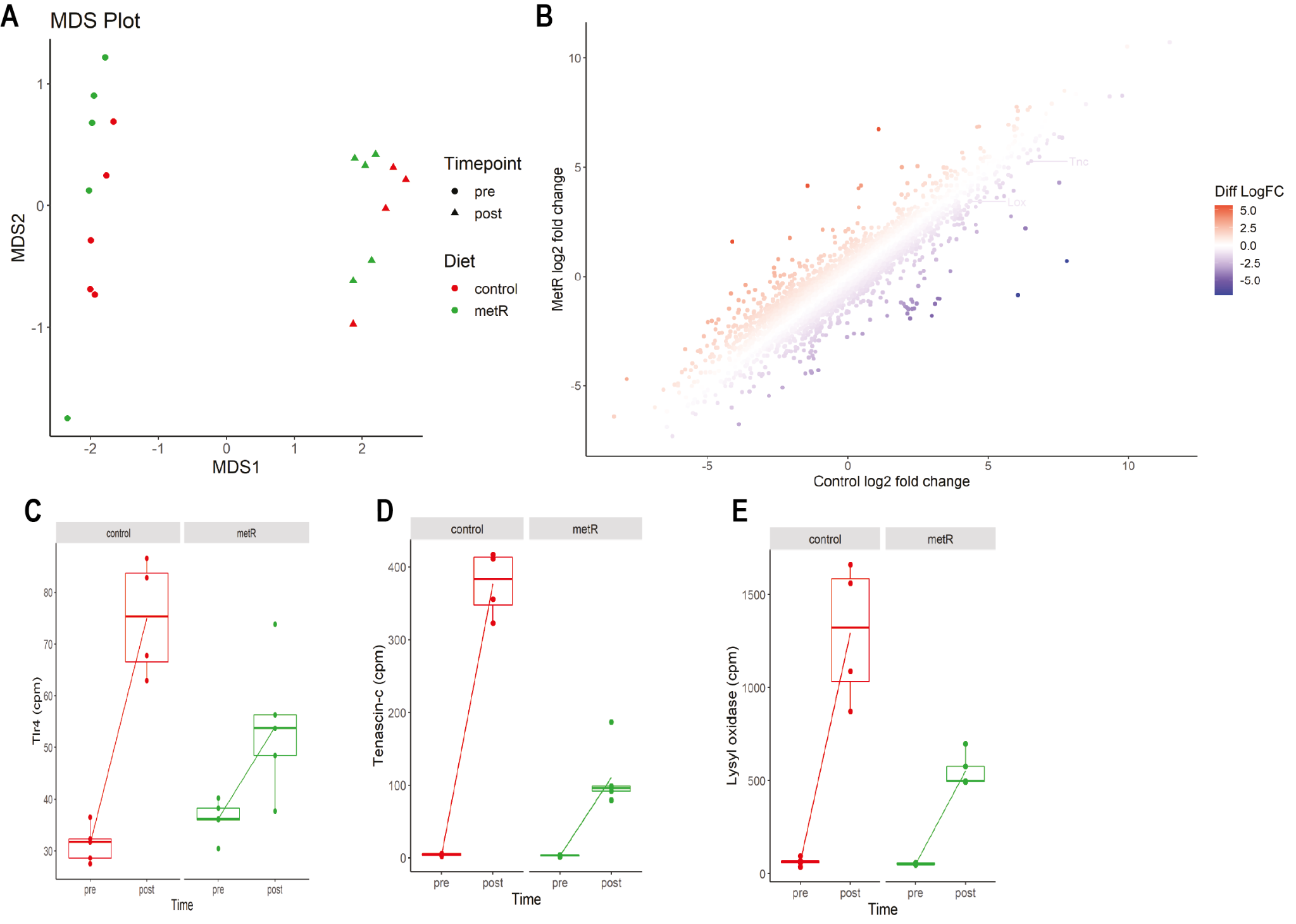

**Supplemental Figure 6.** *Transcriptome analysis of vein graft PVAT at POD1.*
**A:** principal component analysis of pre- and post-surgery PVAT in control-fed and MetR mice. **B:** fold change in transcript expression between caval vein PVAT and vein graft PVAT at POD1 for both control-fed and MetR mice. **C:** expression of tlr4 (in counts per million) in caval vein and vein graft PVAT from control-fed and MetR mice. **D**: tenascin-c transcript expression (in counts per million) in control-fed and MetR mice caval vein and vein graft PVAT. **E**: lysyl oxidase transcript expression (in counts per million) in control-fed and MetR mice caval vein and vein graft PVAT.

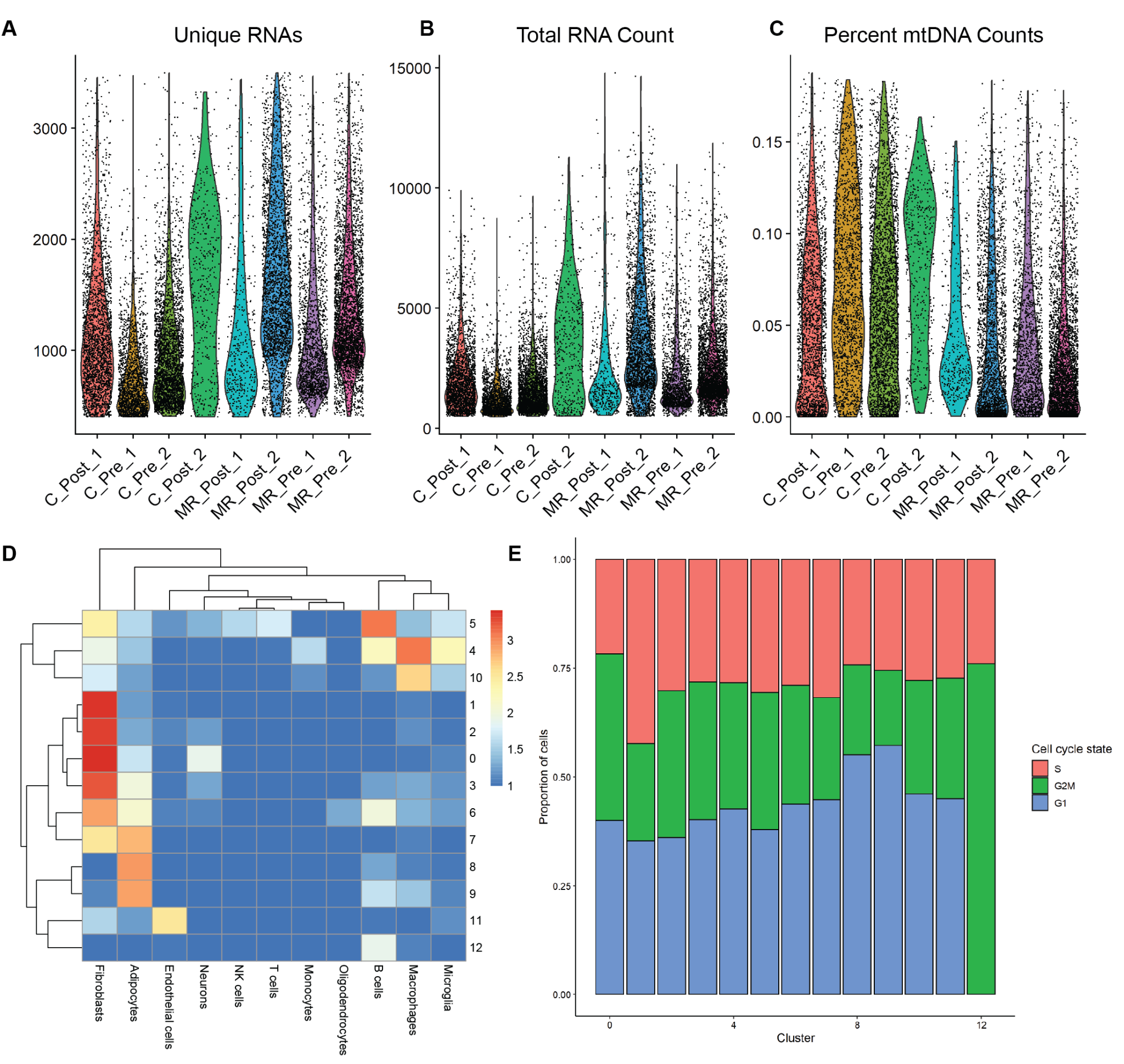

**Supplemental Figure 7.** *Single-cell nuclear sequencing analysis of pre-op and POD1 PVAT.*

**A:** Unique RNAs present per sample. **B:** Total RNA count per sample. **C:** Percent mitochondrial DNA (mtDNA) as percentage of total RNA per sample. **D:** Heatmap comparing transcriptome signature from cell types commonly present in PVAT with our cell clusters (0 – 12). **E:** Cell-cycle state as a proportion of total cells per cluster.

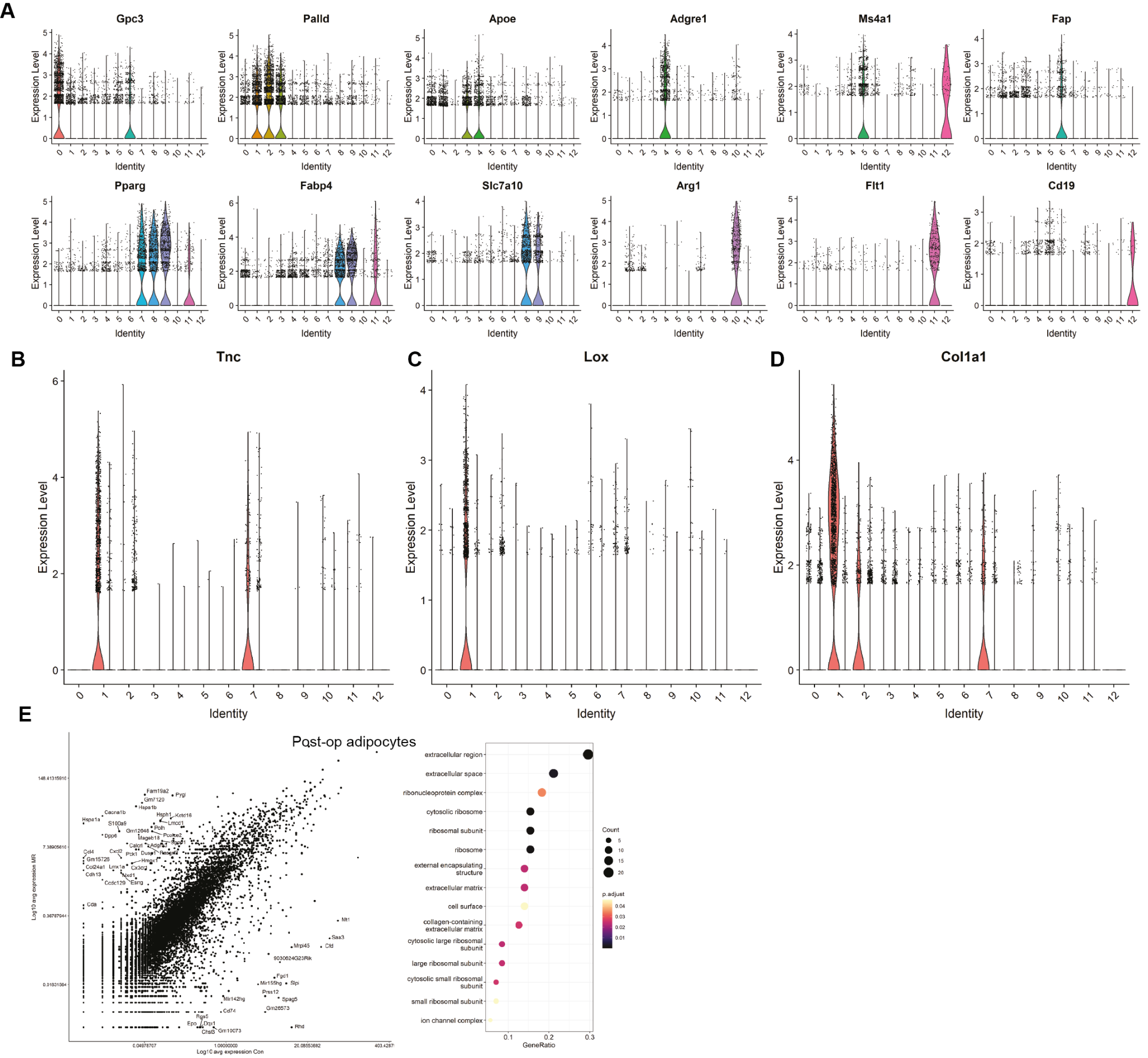

**Supplemental Figure 8.** *Single-cell nuclear sequencing analysis of pre-op and POD1 PVAT*.

***A:*** Validity of selected markers for defining each cell cluster. **B-D:** Expression levels of Tnc, Lox and Col1a1 transcripts in different clusters. **E:** Transcript expression levels and pathway analysis in post-operative adipocytes.

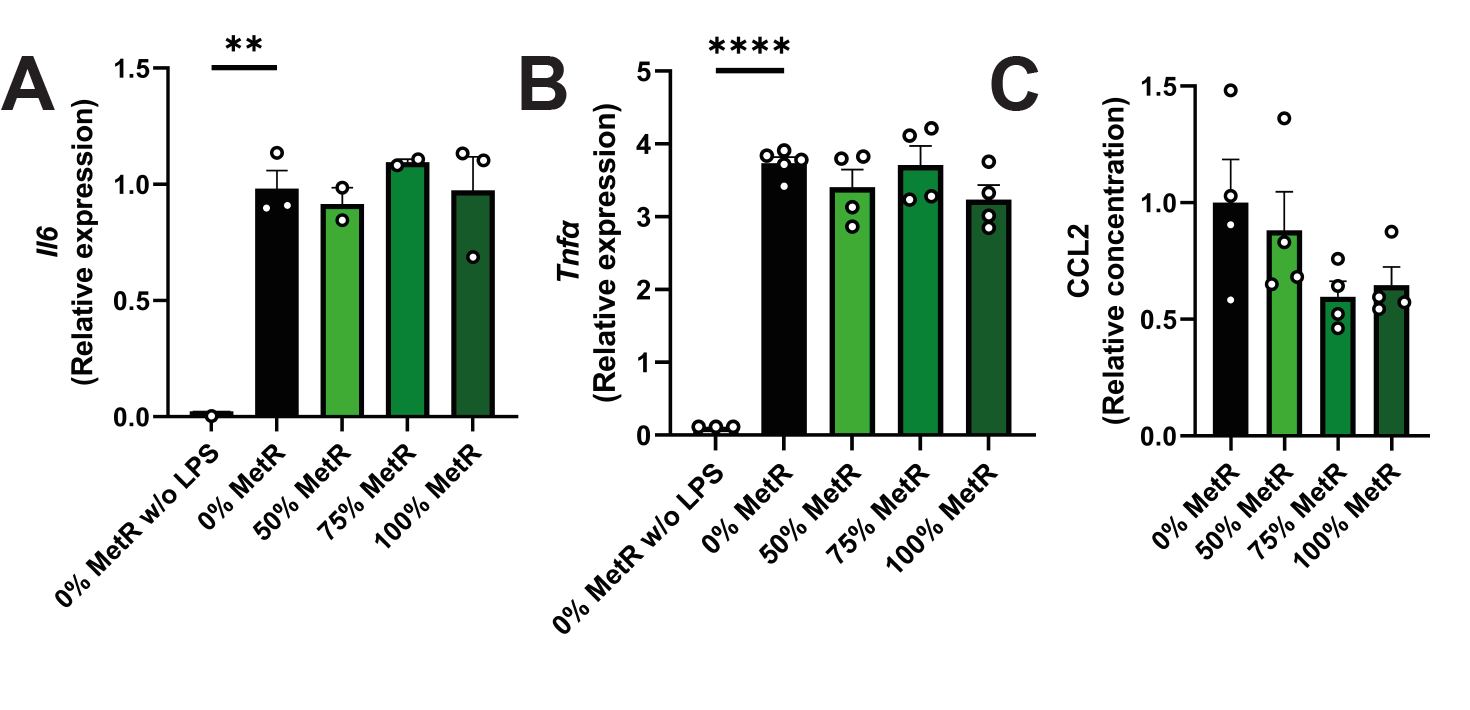

**Supplemental Figure 9. Bone marrow-derived macrophages subjected to MetR in vitro**

**A:** qPCR for *Il6* in pro-inflammatory macrophages under control and MetR conditions. **B:** qPCR for *Tnfα* in pro-inflammatory macrophages under control and MetR conditions. **C:** supernatant from macrophage MetR experiment tested for CCL2 cytokine production via ELISA. Graphs are presented as mean ± SEM. ** P<0.01, ** P<0.0001

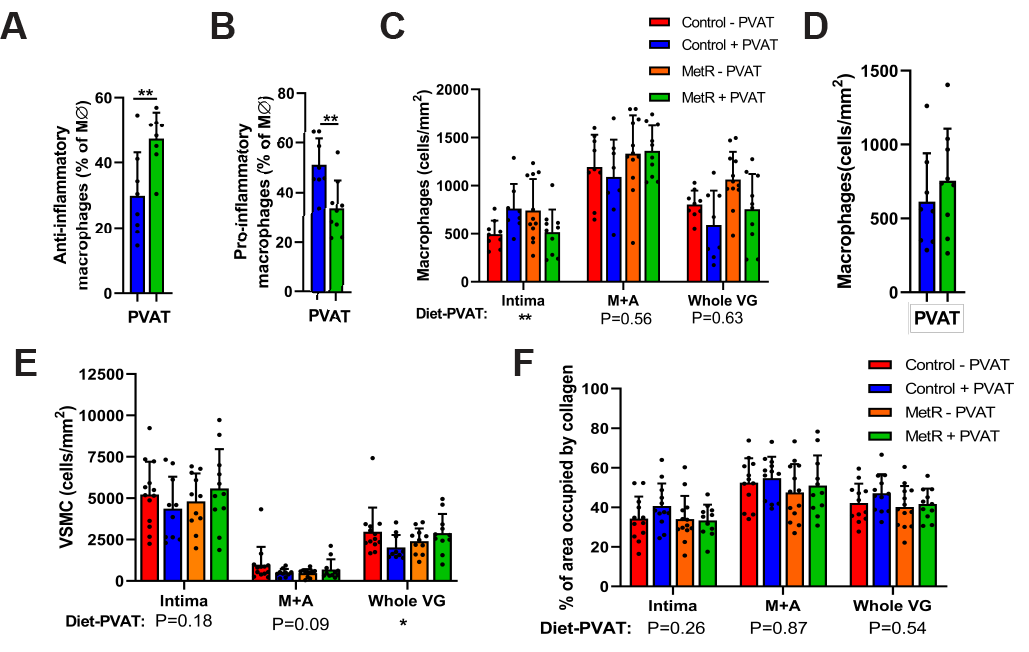

**Supplemental Figure 10.** *VSMC density, collagen content and MΦ macrophages in Vein Grafts and PVAT at POD28.*

**A**: Pro-inflammatory macrophage count in PVAT of control-fed and MetR mice, in cells/mm^2^. **B**: Anti-inflammatory macrophage count in PVAT of control-fed and MetR mice, in cells/mm^2^. C: MΦ-macrophages per vein graft layer in cells/mm^2^. **D**: MΦ-macrophage count in PVAT of control-fed and MetR mice, in cells/mm^2^. **E**: VSMC density per vein graft layer in cells/mm^2^. **F**: percentage of vein graft layer occupied by collagen. All statistical testing was done via two-way ANOVA with Tukey’s multiple comparisons test, unless indicated otherwise, n=10-13/group. ****** P<0.01.

| **Histomorphometric**  **parameter** | **Source of Variation** | **% of total variation** | **P-value** | **P-value summary** |
| --- | --- | --- | --- | --- |
|  | Interaction | 9.146 | 0.0146 | * |
| I/M area ratio | PVAT +/- | 0.006033 | 0.9482 | ns |
| **(Fig. 2F)** | Diet | 31.16 | <0.0001 | **** |
|  | Interaction | 7.538 | 0.0324 | * |
| I/M thickness ratio | PVAT +/- | 0.001153 | 0.9783 | ns |
| (**Fig. 2G**) | Diet | 24.40 | 0.0003 | *** |
|  | Interaction | 4.274 | 0.1451 | ns |
| Intimal area | PVAT +/- | 0.3439 | 0.6760 | ns |
| (**Fig. S2A**) | Diet | 10.96 | 0.0219 | * |
|  | Interaction | 2.284 | 0.2734 | ns |
| Media area | PVAT +/- | 0.4127 | 0.6397 | ns |
| (**Fig. 2I**) | Diet | 16.50 | 0.0047 | ** |
|  | Interaction | 2.507 | 0.2736 | ns |
| Intimal thickness | PVAT +/- | 1.591 | 0.3819 | ns |
| **(Fig. S2B**) | Diet | 6.780 | 0.0751 | ns |
|  | Interaction | 8.223 | 0.0159 | * |
| Media thickness | PVAT +/- | 1.254 | 0.3325 | ns |
| **(Fig. 2H)** | Diet | 35.61 | <0.0001 | **** |
|  | Interaction | 6.514 | 0.0719 | ns |
| Lumen area | PVAT +/- | 0.2124 | 0.7407 | ns |
| **(Fig. S2C)** | Diet | 10.26 | 0.0253 | * |

**Table 1.** Percentage of variation between groups that can be explained by diet, PVAT or a diet-PVAT interaction. Two-way ANOVA with Tukey’s multiple comparison test on histomorphometric parameters in **Figure 2** & **Fig. S2.** n=11-13/group.

| **Parameter** | **Source of Variation** | **% of total variation** | **P-value** | **P-value summary** |  | **Parameter** | **Source of Variation** | **% of total variation** | **P-value** | **P-value summary** |
| --- | --- | --- | --- | --- | --- | --- | --- | --- | --- | --- |
| Pro-/anti-inflammatory macrophage ratio | Interaction | 20.38 | 0.0016 | ** |  | Pro-inflammatory macrophages | Interaction | 10.33 | 0.0489 | * |
| Intima | PVAT +/- | 0.3913 | 0.6384 | ns |  | Intima | PVAT +/- | 0.6856 | 0.6025 | ns |
| **(Fig. 3A)** | Diet | 20.11 | 0.0017 | ** |  | **(Fig. 3E)** | Diet | 1.899 | 0.3876 | ns |
| Pro-/anti-inflammatory macrophage ratio | Interaction | 10.70 | 0.018 | * |  | Pro-inflammatory macrophages | Interaction | 3.346 | 0.2238 | ns |
| Media | PVAT +/- | 1.663 | 0.3345 | ns |  | Media | PVAT +/- | 2.986 | 0.2500 | ns |
| **(Fig. 3A)** | Diet | 28.19 | 0.0003 | *** |  | **(Fig. 3E)** | Diet | 17.65 | 0.0074 | ** |
| Pro-/anti-inflammatory macrophage ratio | Interaction | 19.86 | 0.0006 | *** |  | Pro-inflammatory macrophages | Interaction | 6.346 | 0.1039 | ns |
| Whole VG | PVAT +/- | 0.9584 | 0.4160 | ns |  | Whole VG | PVAT +/- | 2.260 | 0.3259 | ns |
| **(Fig. 3A)** | Diet | 31.89 | <0.0001 | **** |  | **(Fig. 3E)** | Diet | 11.99 | 0.0278 | * |
| Anti-inflammatorymacrophage | Interaction | 12.01 | 0.0173 | * |  | MΦ- macrophages | Interaction | 20.02 | 0.0054 | ** |
| intima | PVAT +/- | 4.668 | 0.1284 | ns |  | intima | PVAT +/- | 0.1291 | 0.8133 | ns |
| **(Fig. 3D)** | Diet | 15.95 | 0.0068 | ** |  | **(Fig. S3F)** | Diet | 0.00009889 | 0.9948 | ns |
| Anti-inflammatory macrophages | Interaction | 4.024 | 0.2121 | ns |  | MΦ- macrophages | Interaction | 0.8676 | 0.5669 | ns |
| Media | PVAT +/- | 1.793 | 0.4019 | ns |  | Media | PVAT +/- | 0.2800 | 0.7446 | ns |
| **(Fig. 3D)** | Diet | 7.059 | 0.1012 | ns |  | **(Fig. S3F)** | Diet | 8.404 | 0.0806 | ns |
| Anti-inflammatory macrophages | Interaction | 8.175 | 0.0594 | ns |  | MΦ- macrophages | Interaction | 0.4932 | 0.6300 | ns |
| Whole VG | PVAT +/- | 2.186 | 0.3205 | ns |  | Whole VG | PVAT +/- | 14.91 | 0.0113 | * |
| **(Fig. 3D)** | Diet | 14.73 | 0.0130 | * |  | **(Fig. S3F)** | Diet | 9.798 | 0.0372 | * |

**Table 2.** Percentage of variation between groups that can be explained by diet, PVAT or a diet-PVAT interaction. Two-way ANOVA with Tukey’s multiple comparison test on pro-/anti-inflammatory macrophage ratio and anti-inflammatory macrophages (left), anti-inflammatory macrophages and MΦ macrophages (right). in **Figure 3** & **Fig. S3.** n=10-13/group.

| **Parameter** | **Source of Variation** | **% of total variation** | **P-value** | **P-value summary** |  | **Parameter** | **Source of Variation** | **% of total variation** | **P-value** | **P-value summary** |
| --- | --- | --- | --- | --- | --- | --- | --- | --- | --- | --- |
| % VSMC of | Interaction | 6.622 | 0.0897 | ns |  | VSMC/mm^2^ | Interaction | 4.246 | 0.1832 | ns |
| Intima | PVAT +/- | 0.1041 | 0.8286 | ns |  | Intima | PVAT +/- | 0.008998 | 0.9506 | ns |
| **(Fig. 3G)** | Diet | 1.583 | 0.4005 | ns |  | **(Fig. S3E)** | Diet | 1.033 | 0.5080 | ns |
| % VSMC of | Interaction | 4.430 | 0.1693 | ns |  | VSMC/mm^2^ | Interaction | 6.810 | 0.0860 | ns |
| Media | PVAT +/- | 0.1700 | 0.7855 | ns |  | Media | PVAT +/- | 1.037 | 0.4962 | ns |
| **(Fig. 3G)** | Diet | 0.1062 | 0.8296 | ns |  | **(Fig. S3E)** | Diet | 1.278 | 0.4504 | ns |
| % VSMC of | Interaction | 13.01 | 0.0153 | * |  | VSMC/mm^2^ | Interaction | 10.20 | 0.0355 | * |
| Whole VG | PVAT +/- | 0.2532 | 0.7262 | ns |  | Whole VG | PVAT +/- | 1.095 | 0.4802 | ns |
| **(Fig. 3G)** | Diet | 2.068 | 0.3194 | ns |  | **(Fig. S3E)** | Diet | 0.4385 | 0.6545 | ns |
| VSMC+ Ki-67 | Interaction | 0.9706 | 0.5219 | ns |  | % collagen of | Interaction | 2.705 | 0.2627 | ns |
| intima | PVAT +/- | 0.01213 | 0.9428 | ns |  | Intima | PVAT +/- | 1.738 | 0.3681 | ns |
| **(Fig. 3H)** | Diet | 1.456 | 0.4334 | ns |  | **(Fig. S3E)** | Diet | 3.099 | 0.2311 | ns |
| VSMC+ Ki-67 | Interaction | 4.180 | 0.1743 | ns |  | % collagen of | Interaction | 0.05245 | 0.8773 | ns |
| Media | PVAT +/- | 3.468 | 0.2151 | ns |  | Media | PVAT +/- | 1.229 | 0.4562 | ns |
| **(Fig. 3H)** | Diet | 0.8681 | 0.5323 | ns |  | **(Fig. S3E)** | Diet | 2.802 | 0.2625 | ns |
| VSMC+ Ki-67 | Interaction | 2.150 | 0.3458 | ns |  | % collagen of | Interaction | 0.7912 | 0.5427 | ns |
| Whole VG | PVAT +/- | 0.5066 | 0.6458 | ns |  | Whole VG | PVAT +/- | 2.748 | 0.2591 | ns |
| **(Fig. 3H)** | Diet | 0.6122 | 0.6135 | ns |  | **(Fig. S3E)** | Diet | 3.815 | 0.1848 | ns |

**Table 3.** Percentage of variation between groups that can be explained by diet, PVAT or a diet-PVAT interaction. Two-way ANOVA with Tukey’s multiple comparison test on either % of vein graft layers occupied by VSMC, VSMC/mm^2^, VSMC colocalizing with Ki-67 or % of vein graft layers occupied by collagen. As depicted in graphs in **Figure 3** & **Fig. S3.** n=10-13/group.
